# Wiring multiple microenvironment proteomes uncovers the biology in head and neck cancer

**DOI:** 10.1101/2021.10.22.465341

**Authors:** Ariane Fidelis Busso-Lopes, César Rivera, Leandro Xavier Neves, Daniela Campos Granato, Fábio Malta de Sá Patroni, Tatiane de Rossi Mazo, Ana Gabriela Costa Normando, Romênia Ramos Domingues, Henry Heberle, Marco Antônio Pretti, Barbara Pereira de Mello, Andre Nimtz Rodrigues, Pammela Araujo Lacerda, Nayane Alves de Lima Galdino, Kenneth John Gollob, Tiago da Silva Medina, Nilva de Karla Cervigne, Ana Carolina Prado-Ribeiro, Thaís Bianca Brandão, Luisa Lina Villa, Miyuki Uno, Mariana Boroni, Luiz Paulo Kowalski, Wilfredo González-Arriagada, Adriana Franco Paes Leme

## Abstract

The poor prognosis of head and neck cancer (HNC) is associated with the presence of metastasis within the lymph nodes (LNs). Herein, the proteome of 140 multisite samples from a 59-HNC patient cohort, including primary and matched LN-negative or -positive tissues, saliva, and blood cells, reveals insights into the biology and potential metastasis biomarkers that may assist in clinical decision making. Protein profiles are strictly associated with immune modulation across datasets, and this provides the basis for investigating immune markers associated with metastasis. The proteome of LN metastatic cells recapitulates the proteome of the primary tumor sites. Conversely, the LN microenvironment proteome highlights the candidate prognostic markers. By integrating prioritized peptide, protein, and transcript levels with machine learning models, we identified a nodal metastasis signature in the blood and saliva. In summary, we present the deepest proteome characterization wiring multiple sampling sites in HNC, thus providing a promising basis for understanding tumoral biology and identifying metastasis-associated signatures.

## INTRODUCTION

Head and neck cancer is the eighth leading cause of cancer worldwide, and 90% of these tumors are mucosal head and neck squamous cell carcinomas (HNSCC) (Sung et al., 2021). HNSCC can severely impact the quality of life of patients due to treatment sequelae and high rates of locoregional recurrences. Unfavorable outcomes are largely related to the presence of lymph node metastasis that reduces survival by approximately 50%, and this is the primary argument supporting the use of elective neck treatment (Ho et al., 2017). The detection of lymph node alterations can be challenging, and the discovery of molecular markers that can allow for an accurate identification of locoregional spread would enable clinicians to avoid unnecessary extensive operations and could reduce postoperative morbidity (Kowalski and Sanabria, 2007).

While cancer research has previously focused on characterizing malignant cells in primary tumor tissues, the investigation of additional environments implicated in HNSCC regulation may lead to an improved understanding of the mechanisms underlying carcinogenesis (Carnielli et al., 2018; Puram et al., 2017). In addition to cancer cells, the tumor microenvironment (TME) comprises distinct cell subsets, as the immune portion and cancer-associated fibroblasts (CAFs), and it is of special interest once the intense crosstalk among these heterogeneous populations reprogram key processes responsible for tumor growth and invasion (Hanahan and Coussens, 2012; Hanahan and Weinberg, 2011). Additionally, neoplastic cells can enter lymphatic vessels and migrate to lymph nodes where they interact with the host immune environment and establish metastasis (Hoshida et al., 2006; Jones et al., 2018). Thus, a better understanding of the molecular signals within the TME and in the metastatic microenvironment may provide insights into tumor biology, thus helping to guide clinical investigations.

The composition of body fluids, that is able to wire diverse microenvironments, can also be affected by cancer. Tumor-specific T cells, circulating tumor cells (CTCs), macrophage-like cells, tumor endothelial cells, cancer-associated fibroblasts (CAFs), free molecules, and exosomes have all been identified in the peripheral blood of cancer patients and are valuable in clinical decision-making (Adams et al., 2014; Ao et al., 2015; Cima et al., 2016; Cohen et al., 2015; Niccolai et al., 2016). Saliva has also been proven to be a promising source of biomarkers in HNSCC due to its proximity to tumor lesions (Neves et al., 2020; Wang et al., 2015; Winck et al., 2015). Thus, the analysis of fluids or liquid biopsies raises the possibility of probing the molecular profile of tumors in a non-invasive manner and may provide a valuable means of tracking biomarkers in HNSCC.

In this scenario, clinical proteomics has emerged as a promising approach for the identification and quantification of potential markers, leveraging the development of new tools that can be used in clinical practice. Technological advances in proteomics, particularly in the mass spectrometry (MS) field (Yates, 2019), have the power to provide a deeper understanding of the molecular mechanisms and guide the discovery of biomarkers (Huang et al., 2021; Irmisch et al., 2021; Sinha et al., 2019; Uzozie and Aebersold, 2018). Herein, we used a multisite mass spectrometry-based discovery approach in a 59-patient cohort, and this was followed by a deep biological characterization of the proteomes and application of a multiparametric machine learning model to prioritize targeted molecules. Taken together, this study presents the basis for understanding the response of multiple microenvironments to lymph node metastasis and indicates prognostic signatures in HNSCC.

## RESULTS

### Global proteomes are collectively implicated in translation and the immune response

To obtain a comprehensive view of the proteome composition in HNSCC, we selected 27 primary tumors, 27 matched negative or positive lymph nodes, 24 buffy coats, and 24 saliva samples from a 59-patient cohort to evaluate the protein content using label-free quantitative mass spectrometry in the discovery phase **(Table S1)**. This group of samples included 27 matched formalin-fixed paraffin-embedded (FFPE) tumors and lymph node tissues and also 15 paired buffy coat and saliva samples. A histology-guided approach was employed to harvest malignant and non-malignant enriched cell populations in both primary tumor and lymph node tissues. While malignant cells refer to the tumoral or metastatic cells themselves, the non-malignant portion includes microenvironment cells that surround and support the malignant populations (Anderson and Simon, 2020). The populations that were evaluated included (i) malignant cells from primary tumors, (ii) non-malignant cells located adjacent to primary tumors (mucosal margins), (iii) malignant cells from metastatic lymph nodes, (iv) non-malignant cells located adjacent to sites of metastasis from lymph nodes, (v) buffy coat samples, and (vi) saliva cell samples. Mass spectrometry quality control measures were utilized for all the experiments **(Figure S1)**. Two primary tumor samples (malignant cells) were excluded due to inconsistent detection of control peptide precursor ions (patients 2875 and 4417), and this resulted in 25 remaining malignant samples from primary tumors that were used for analysis. We identified an average of 2,048 protein groups that exhibited MS signals covering close to five orders of magnitude **(Figure 1A; Table S2-1 to 7)**. Malignant cells from primary sites yielded the highest number of identified proteins (n = 2,451 proteins), and this was followed by malignant cells from lymph nodes (n = 2,374 proteins). The ranking according to MS signal revealed a buffy coat proteome with a wider dynamic range among all of the sites evaluated (n = 2,188 proteins), although different LC gradients were used. A total of 313 proteins were shared across multiple sites **(Figure S2A)**.

**Figure 1.**
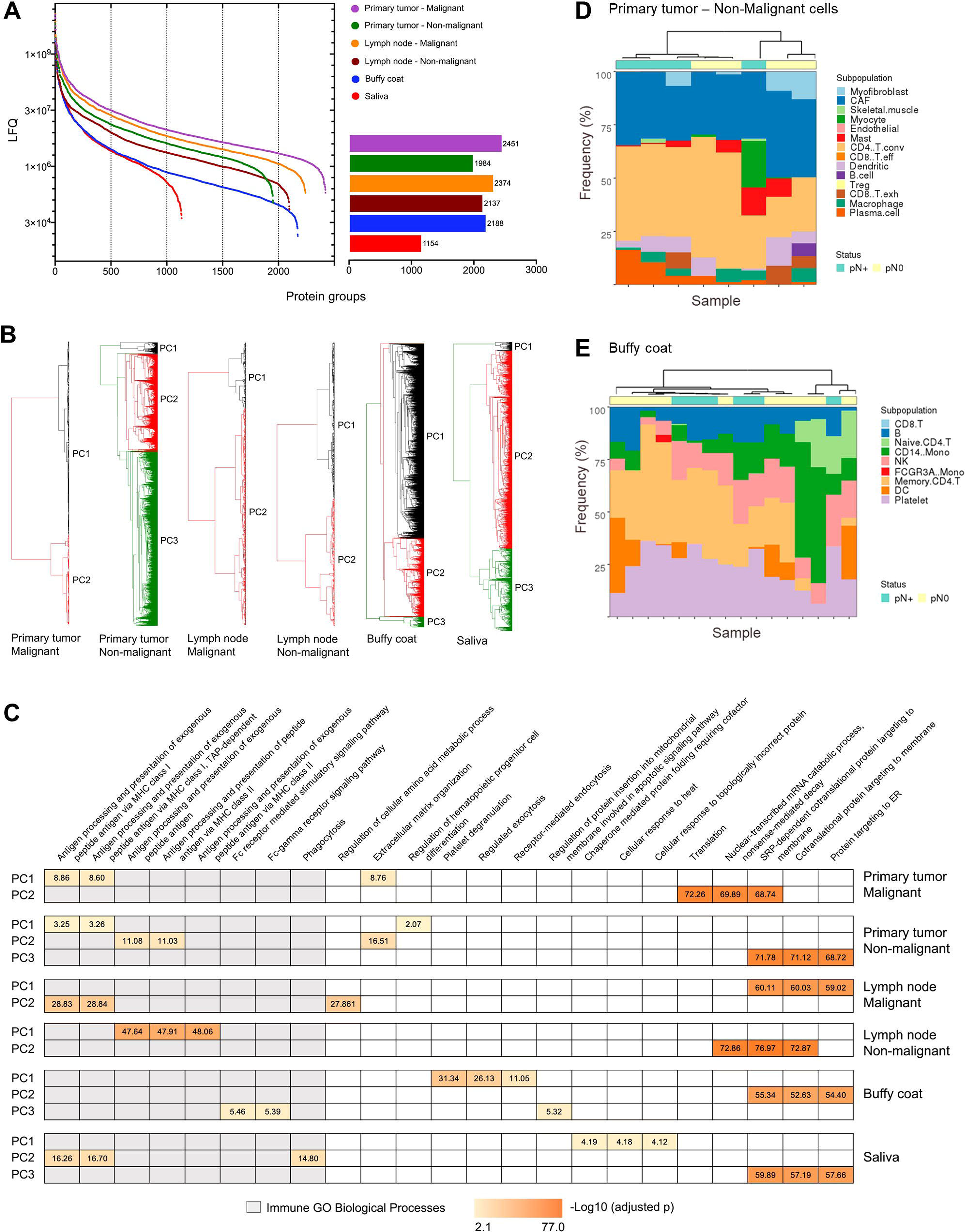
Proteomic profile of tissues and fluids in a 59-HNSCC patient cohort. (A) Dynamic range of proteomics quantitative data for primary tumor malignant (n = 25 samples) and non-malignant (n = 27 samples) cells, lymph node malignant (n = 13 samples) and non-malignant (n = 27 samples) cells, and buffy coat (n= 24 samples) and saliva samples (n = 24 samples). (B) Groups identified by clustering of the protein datasets for the multisites. Grouping was generated using Ward Sqeuclidean (primary tumor – malignant: 2,451 proteins), Weighted Camberra (primary tumor – non-malignant: 1,984 proteins and buffy coat: 2,188 proteins), Ward Sqeuclidean (lymph node – malignant: 2,374 proteins and lymph node – non-malignant: 2,137 proteins), and Complete Sqeuclidean (Saliva: 1,154 proteins). (C) Top-3 significant GO biological processes enriched for the PC groups (adjusted p ≤ 0.05). (D-E) Predicted composition of immune populations in non-malignant cells from primary tumors (D) and buffy coat (E) samples using public databases and CIBERSORTx. Each bar represents a sample. Samples were clustered using the Ward.D distance, and the pN status was annotated. See also Figures S1-S3.

We then investigated the HNSCC proteome to retrieve insights from the biology of multiple sites. Proteins from tissues and fluids were separated into clusters based on the hierarchical relationship among the label-free quantitation (LFQ) intensities (PC: protein cluster) **(Figure 1B; Figure S2B)**, and the PCs were associated with Gene Ontology (GO) biological processes (adjusted p ≤ 0.05) **(Figure 1C)**. Translational processes were enriched in PCs from all HNSCC sites with high significance (−Log_10_ [adjusted p] = 52.63 to 76.97). At least one PC from each region was enriched for immune-related processes that specifically included antigen processing and presentation via MHC class I in malignant cells from primary tumors and lymph nodes, non-malignant cells adjacent to the tumor, and saliva samples (GO:0042590/GO:0002479; −Log_10_ [adjusted p] = 3.25 to 28.84), antigen processing and presentation via MHC class II for non-malignant cells adjacent to the tumor and lymph node sites (GO:0002495/GO:0019886; −Log_10_ [adjusted p] = 11.03 to 48.06), Fc receptor signaling pathways in buffy coat samples (GO:0002431/ GO:0038094; −Log_10_ [adjusted p] = 5.39 to 5.46), and phagocytosis in saliva samples (GO:0006909; −Log_10_ [adjusted p] = 14.80).

Taken together, these findings revealed specific protein composition for malignant and non-malignant cells derived from primary tumors and lymph nodes, buffy coat, and saliva samples. We also identified subsets of proteins across distinct cell populations that exhibit similar abundance profiles and may modulate common biological functions in HNSCC, particularly immune response and translation-related processes.

### Protein profiles indicate specific immune phenotypes across datasets

Based on the observation that proteomes from multiple sites were enriched for immune processes **(Figure 1C)**, we inferred the immune composition associated with our bulk proteomes based on signatures from publicly available single-cell RNA sequencing data. Non-immune subpopulations were also inferred from the proteomic data.

First, the global proteome levels from primary tumors (malignant cells) were compared to the transcriptomic levels using data from 500 HNSCC tumor samples retrieved from The Cancer Genome Atlas (TCGA). We observed a moderate degree of correlation between proteome and transcriptome levels (ρ = 0.53; p ≤ 0.05), thus indicating that RNA-based cell signatures could be used to estimate subpopulations from bulk proteomes **(Figure S3A)**. Subsequently, public scRNASeq datasets comprised of 18 HNSCC tissue samples (Puram et al., 2017) and a human peripheral blood mononuclear cell (PBMC) sample from a healthy donor (https://www.10xgenomics.com) were used to generate a signature of microenvironment (tumor and lymph node) and blood samples. This matrix was applied as a reference to deconvolute whole proteome information from non-malignant cells and fluids, respectively, in CIBERSORTx (Newman et al., 2019) **(Figure S3B-C)**. According to the prediction, non-malignant cells from tumors were enriched with high fractions of CD4+ T cells (eight out of eight samples; deconvolution p ≤ 0.1) **(Figure 1D)**. Mast cells, dendritic cells, macrophages, and plasma cells were frequently detected in HNSCC (six to seven out of eight samples; deconvolution p ≤ 0.1); however, they were present at lower percentages. It is of interest to highlight the enrichment of the non-immune population of CAFs in all 8 HNSCC tissue samples, as this observation indicates that further studies focusing on CAFs may benefit these patients. Remarkably, the predominance of T lymphocytes and fibroblasts in HNSCC tumors has been demonstrated previously at the RNA level and agrees with predictions based on the use of tumor proteomes (Puram et al., 2017). Buffy coat prediction revealed a high percentage of memory CD4+ T cells, CD14 monocytes, B cells, and NK cells (14 to 15 out of 16 samples; deconvolution p ≤ 0.1) **(Figure 1E)**, thus revealing that distinct immune profiles could be inferred for the tumor microenvironment and blood cells. The presence (pN+) or absence (pN0) of nodal metastasis could not segregate samples perfectly using hierarchical clustering **(Figure 1D-E)**. The immune and non-immune subpopulations could not be inferred for saliva cells due to a lack of a suitable scRNASeq reference and for lymph node non-malignant samples due to non-significant results (deconvolution p > 0.01). These analyses represent a detailed immune characterization of the proteome from HNSCC microenvironments and can be used to predict immune populations enriched in multiple sites based on proteomic data.

### HNSCC multi-sites exhibit immune-associated nodal metastasis markers

We further explored multiple sites to identify common metastasis-dependent markers (primary tumor – malignant: 11 pN+, 14 pN0 patients; primary tumor and lymph node – non-malignant: 13 pN+, 14 pN0; buffy coat: 11 pN+, 13 pN0; saliva cells: 13 pN+, 11 pN0). Malignant cells from the lymph nodes (pN+) were not included in the analysis, as they did not possess the pN0 counterpart to allow for comparisons. A mean of 106 ± 56 differentially abundant proteins was associated with locoregional metastasis across the tissues and fluids (pN+ vs. pN0; p ≤ 0.05; Student’s t-test or proteins detected exclusively in one group) **(Figure 2A-B; Table S3-1 to 5)**. The highest number of differentially abundant proteins was observed in non-malignant cells from lymph nodes (n = 201 proteins), and this was followed by malignant cells from the primary tumor (n = 110 proteins) and non-malignant cells from the primary tumor samples (n = 85 proteins) **(Figure 2A)**. Additionally, 80 and 54 proteins were associated with nodal status in the buffy coat and saliva samples, respectively **(Figure 2A)**. Remarkably, malignant cells from primary tumor, buffy coat and saliva shared similar frequencies and a certain balance between up- and downregulated proteins, whilst both non-malignant environments showed higher frequencies of proteins with higher abundance in the pN+ condition, indicating that overall protein function is upregulated in these microenvironments **(Figure 2B)**. A comparative GO biological process enrichment for the lymph node metastasis proteins highlighted over-represented immune-related terms at multiple sites and were predominantly associated with granulocyte activation (GO:0036230; three datasets; FDR = 2.10E-12 to 7.17E-07), leukocytes (GO:0002366, GO:0002443; three datasets; FDR = 7.62E-14 to 6.05E-05), myeloid cells (GO:0002275; 3 datasets; FDR = 3.78E-14 to 5.14E-07), and neutrophils (GO:0002283, GO:0042119, GO:0002446, GO:0043312; three and four datasets; FDR = 7.62E-14 to 6.05E-05) **(Figure 2C)**. It is becoming increasingly clear that neutrophils possess various functions that dynamically regulate the metastatic cascade, including a role in establishing a premetastatic niche (Kowanetz et al., 2010; Wu et al., 2015) or via neutrophil extracellular traps (NETs) by mediating the trapping of circulating cancer cells (Cools-Lartigue et al., 2013), or still awakening dormant tumor cells (Albrengues et al., 2018). Our enrichment analysis indicated that neutrophils may be important players in the development of local metastasis in HNSCC. Additionally, a common signature of 23 proteins was significantly associated with lymph node metastasis at two or more sites **(Figure 2D).** The 23 proteins are implicated in a series of immune-associated GO biological processes (FDR ≤ 0.05) **(Figure 2E)** that have already been highlighted in the global and metastasis-dependent profiles **(Figure 1C; Figure 2C),** and this strengthens the relevance of the immune response in HNSCC multisites. In parallel, some of these proteins are associated with metabolism (FDR ≤ 0.05) and reflect the results previously shown in HNSCC metastatic cell-derived extracellular vesicles (Busso-Lopes et al., 2021). Alterations in the metabolic profile have been also demonstrated to be associated with the metastatic potential of cancer cells (Fares et al., 2020).

**Figure 2.**
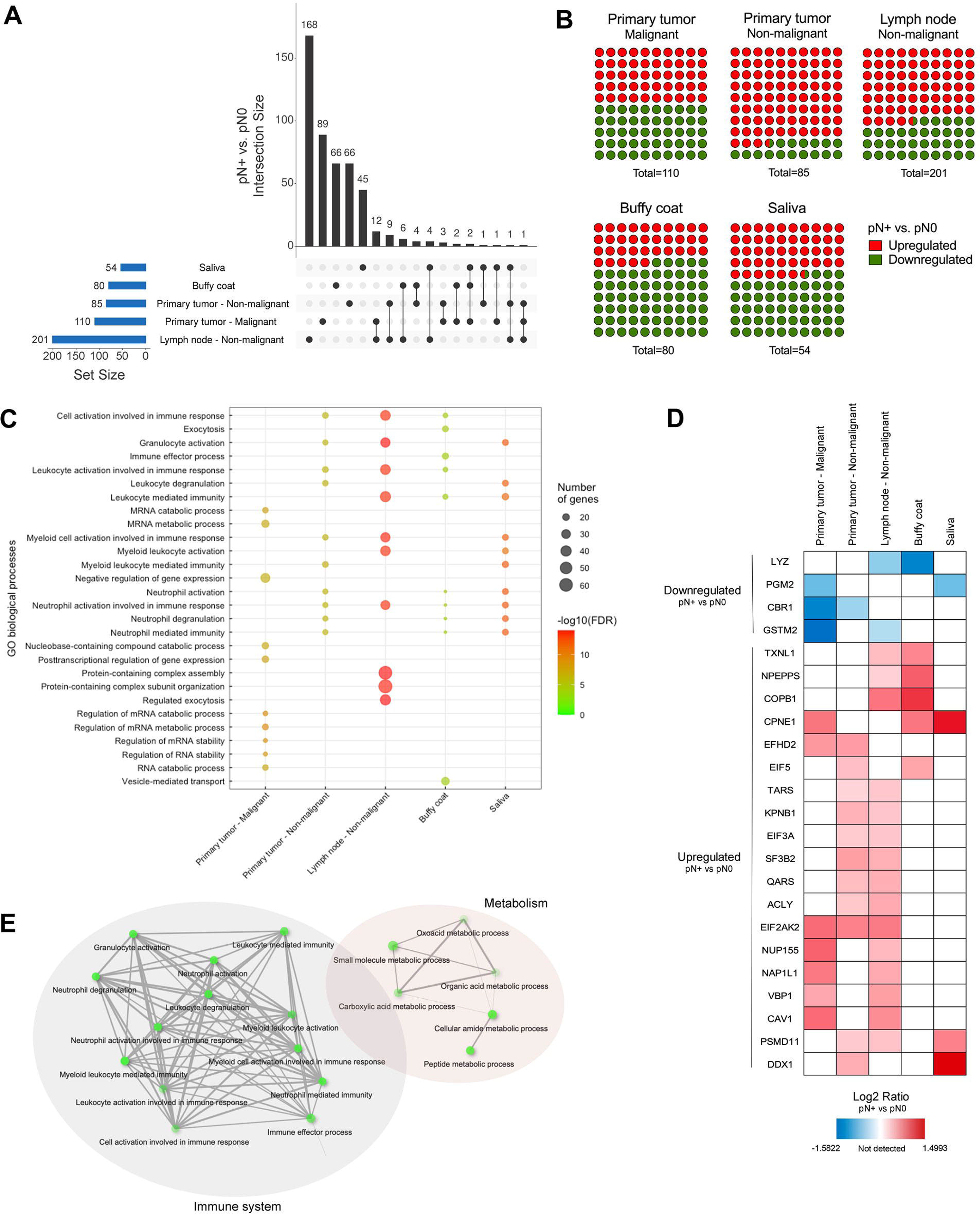
Protein profile associated with lymph node metastasis. (A) Upset plot presenting the intersection of differentially abundant proteins associated with lymph node metastasis in multiple sites (pN+ vs. pN0; p ≤ 0.05; Student’s t-test; primary tumor – malignant: 11 pN+, 14 pN0 patients; primary tumor and lymph node – non-malignant: 13 pN+, 14 pN0; buffy coat: 11 pN+, 13 pN0; saliva cells: 13 pN+, 11 pN0). (B) Frequency of downregulated or upregulated proteins associated with lymph node metastasis in each of the sites. (C) Combined view of the top-10 GO biological processes overrepresented for tissues and fluids considering differentially abundant proteins between pN+ and pN0 from (A) and (B) (Enrichment FDR ≤ 0.01). (D) Abundance of 23 proteins significantly associated with lymph node metastasis in multiple sites (pN+ vs. pN0; p ≤ 0.05; Student’s t-test). (E) GO biological processes significantly enriched for the 23 common proteins listed in (D) (Enrichment FDR ≤ 0.05).

Based on the significance of the immune system in lymph node metastasis, we next investigated if the differentially abundant proteins in the proteomes are cluster markers of immune subpopulations. Cluster markers are herein defined as markers that designate populations via differential expression (**STAR Methods)**. Briefly, cluster markers were identified across HNSCC tissues (Puram et al., 2017) and PBMC scRNASeq public data and compared to differentially abundant proteins (pN+ vs. pN0) identified from our datasets **(Table S3-1 to 5)**. Transcripts from a subset of metastasis-related markers were identified as cluster markers of immune subpopulations, and these included 31 genes from a 201-protein signature from lymph node non-malignant cells that were differentially expressed primarily in CD4+ T cells, plasma cells, and macrophages **(Figure 3A)**, 5/85 genes from primary tumor non-malignant cells that are cluster markers of dendritic cells, mast cells, and CD8+ exhausted T cells **(Figure 3B)**, and 26/80 molecules from the buffy coat that were primarily identified in CD14+ and FCGR3A+ monocytes **(Figure 3C)**. Remarkably, genes such as *ERLIN1, MT1M, VAT1, FBN1, MMP2, STAT1, TOM1* and *SERPINH1* that were identified as potential metastasis markers in the primary tumor or lymph node microenvironments are cluster markers that are exclusively expressed by CAF populations, thus indicating that CAFs can provide an excellent source of candidates as metastasis biomarkers for HNSCC **(Figure 3A-B)**.

**Figure 3.**
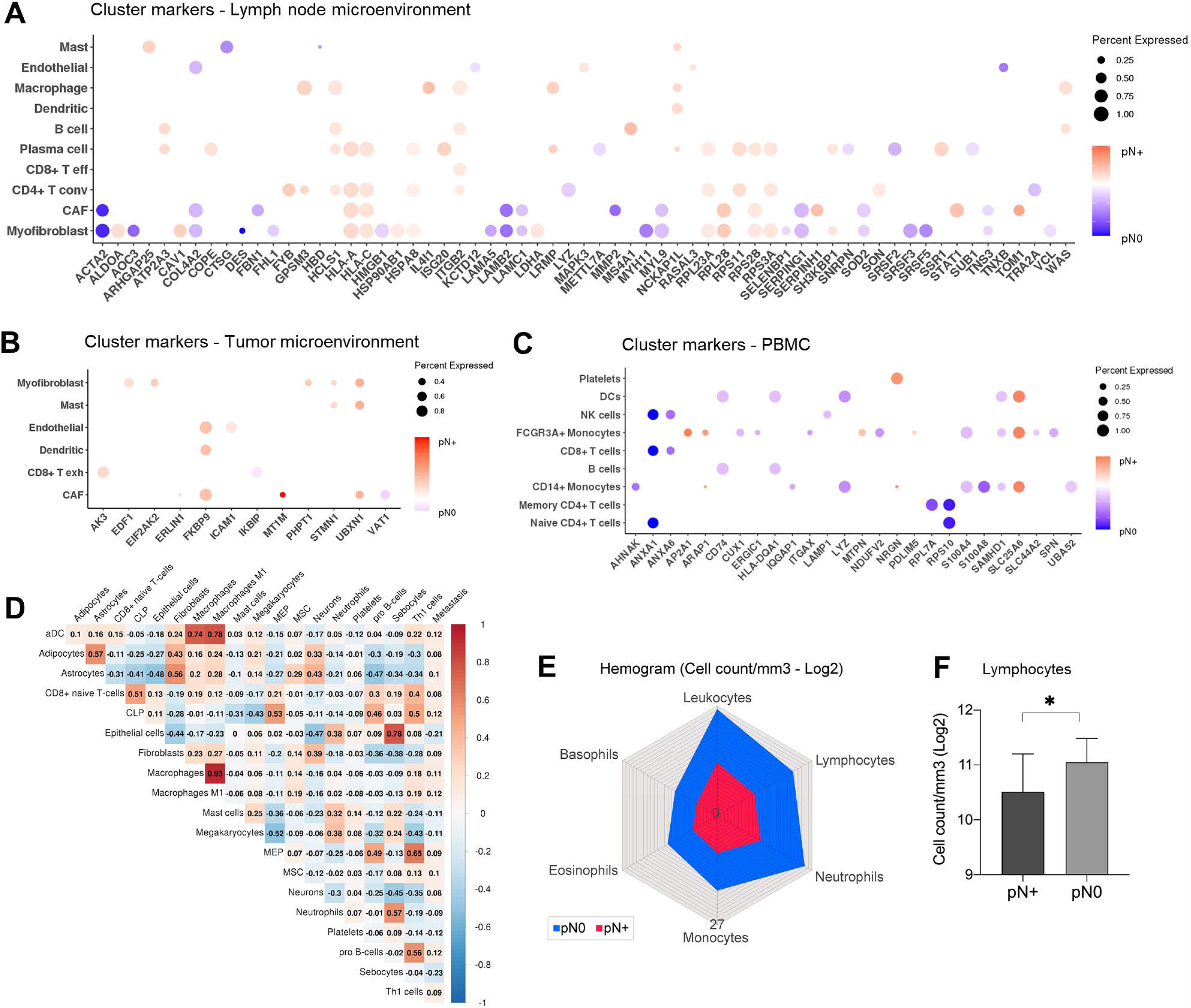
Immune characterization of lymph node metastasis markers using scRNASeq databases. (A-C) Differentially abundant proteins from proteomics data were identified as cluster markers of cells from the lymph node microenvironment (Puram et al., 2017) (A), the tumor microenvironment (Puram et al., 2017) (B), and PBMC (https://www.10xgenomics.com) (C). Colors represent higher LFQ levels in pN+ (red) or pN0 (blue). (D) Spearman correlation coefficients generated for the comparison of nodal status (last column) and immune populations using RNASeq data from TCGA. Correlations with p ≤ 0.05 are presented. (E) Immune cell count/mm^3^ from hemograms collected from 11 pN+ and 14 pN0 HNSCC patients. (F) Association of lymphocyte cell count/mm^3^ with lymph node metastasis (pN+ vs. pN0; p ≤ 0.05; Student’s t-test). *p ≤ 0.05. See also Figure S3.

Next, to reinforce the association between the immune response and lymph node metastasis, we correlated the immune subpopulations that were estimated from bulk RNA sequencing (RNASeq) immune data retrieved from TCGA for a 428-HNSCC (180 pN0, 248 pN+) and a 233-oral squamous cell carcinoma (OSCC; 100 pN0, 133 pN+) cohort with pN outcome. In this context, cell subpopulations were estimated using the in xCell algorithm (Aran, Hu, and Butte, 2017). Interestingly, specific immune populations were correlated with the presence of lymph node metastasis (p ≤ 0.05; Spearman correlation) **(Figure 3D; Figure S3D)**. As a proof-of-concept, we also evaluated in the clinical data from this study cell counts obtained from the clinical laboratory in a group of 25 HNSCC patients (11 pN+, 14 pN0), and our results revealed a lower cell count/mm^3^ of lymphocytes in pN+ compared to that in pN0 (p = 0.0253; Student’s t-test) **(Figure 3E-F)**.

Therefore, through wiring multiple comparisons among our proteomics data, publicly available datasets and cell counting, we strengthened the relevance of the immune system in HNSCC and identified an immune landscape that was associated with the metastatic profile and populations that potentially express metastasis markers. Based on the relevance of the granulocytes/neutrophils, lymphocytes, dendritic cells, monocytes, and macrophages depicted in the biological process enrichment and public data comparison, we prioritized a list of immune markers based on their identification in the proteomes. From the initially 19 selected targets that represent these specific immune populations, nine potential markers passed the quality control assessment and were verified using targeted approaches and further combined with other targets to investigate the ability to distinguish nodal status using machine learning (ML). These immune markers included CD3, CD4, CD8, CD11b, CD14, CD16, CD19, CD45, and CD66b (please see **STAR Methods)**.

### Nodal metastasis cells resemble the molecular signature from tumors

We next sought to explore the protein profile of the malignant and non-malignant populations in the tumor and lymph nodes to elucidate the functional effect of tumor cell spread from the primary site to the lymph nodes. To achieve this, we compared the proteome composition of malignant and non-malignant cells isolated from 11 and 27 matched primary tumors and lymph nodes, respectively.

Of the 2,478 protein groups identified for the malignant portion, 126 proteins were differentially abundant between lymph nodes and primary tumors (q ≤ 0.05; Benjamini-Hochberg test or proteins detected exclusively in one group) **(Table S2-8; Table S3-6)**, thus demonstrating that metastasis recapitulates the content of primary tumors. The differential proteome is strongly involved in actin-based cell movement (Enrichment FDR = 1.22E-04 to 1.79E-02) **(Figure 4A)** that is critical for establishing the epithelial–mesenchymal transition (EMT) necessary for tumor invasion and metastatic dissemination (Savagner, 2001). During EMT, as previously described in HNSCC at the RNA level (Puram et al., 2017), tumor cells undergo a loss of intrinsic polarity and lose cell–cell junctions through extensive reorganization of the cytoskeleton and initiation of actin-based cell motility (Savagner, 2001). Six out of seven proteins enriched for actin-related locomotion were downregulated in the metastatic malignant samples, and these primarily included members of the actin and myosin family (ACTA1, ACTN2, MYH7, MYL1, NEB, and TPM3) **(Figure 4A)**. This likely reduction in cell motility in lymph node metastatic sites compared to that in the primary tumor samples (i) may indicate that distant metastasis predominantly arises independently of lymph node metastasis in HNSCC, as this is known to occur in colorectal cancer where only 35% of cells disseminated to distant sites are related to the ones from locoregional metastasis (Naxerova et al., 2017; Zhang et al., 2020), albeit it is not known in HNSCC. Additionally, this reduction (ii) may be related to the acquisition of a mesenchymal-epithelial transition (MET) program in the metastatic site (the reverse transition of EMT) where metastatic cells recapitulate the pathology and became even less dedifferentiated than are their corresponding primary tumors (Yao et al., 2011) or (iii) may reflect the short follow-up period of the 11 patients included in the analysis (follow-up from 4 to 84 months, mean of 29 months), as most of the HNSCC patients will develop distant metastasis at least 2 years after the initial treatment (Kowalski et al., 2005). None of the 11 patients evaluated in this study possessed distant metastases.

**Figure 4.**
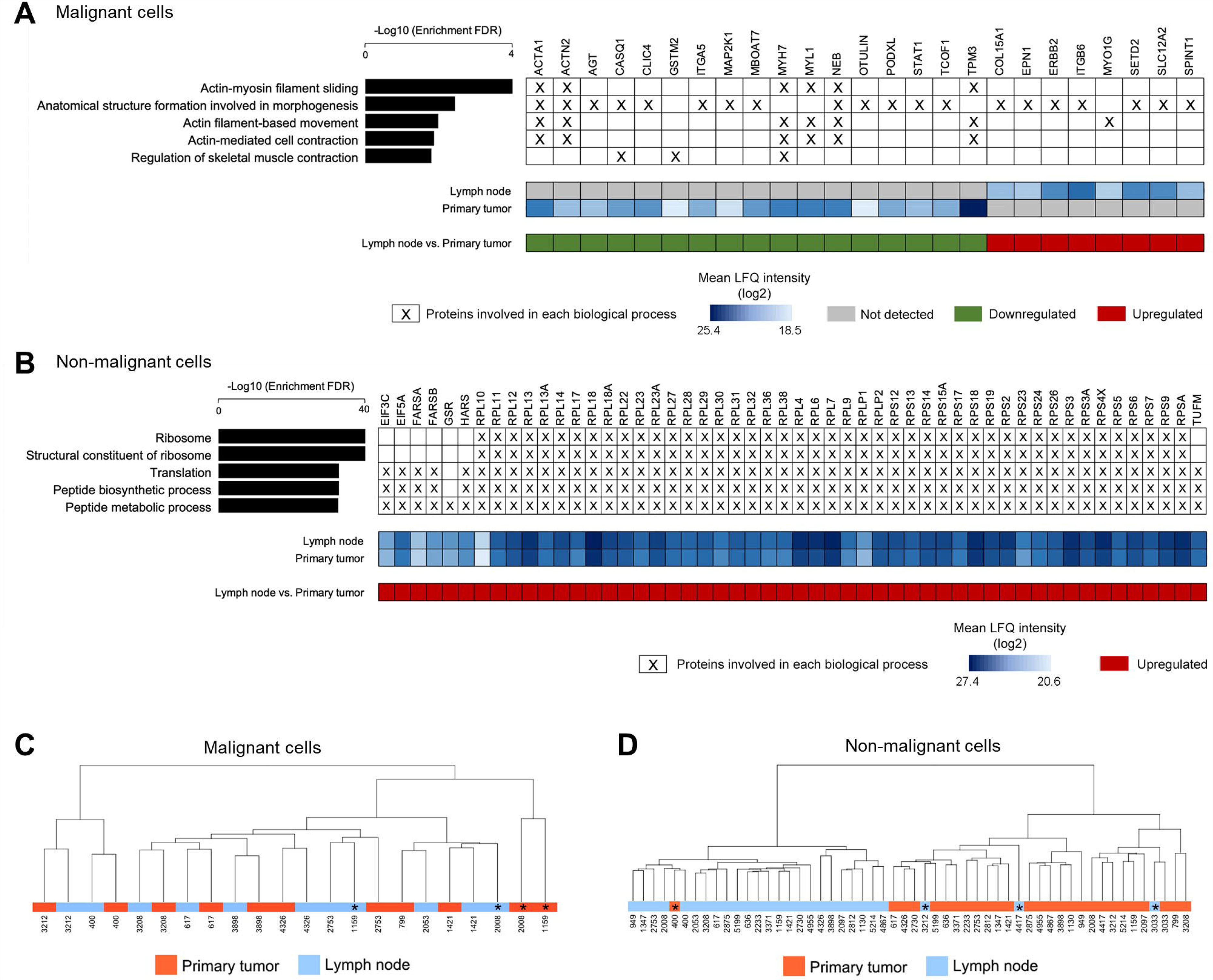
Comparative proteomics analysis of malignant and non-malignant cells from primary tumor and lymph node sites. (A-B) Top-5 GO biological processes enriched for proteins that were differentially abundant between lymph nodes and primary sites identified in malignant (A) and non-malignant (B) samples (Enrichment FDR ≤ 0.05). Proteins involved in each of the top-5 GO enriched biological processes are labeled with an X. Protein abundance in primary sites and lymph nodes are also presented. (C-D) Hierarchical clustering when considering the proteomic profile of malignant (C) and non-malignant (D) cells from primary tumor and lymph nodes (n = 2,478 and 2,360 proteins, respectively). Clustering was performed using the Ward method with Euclidean distance. *Samples that do not follow the main clustering pattern and where the grouping distribution was not associated with any histological or clinical characteristic.

When examining the microenvironment cells (non-malignant cells), 2,360 protein groups were identified, and 873 of these were differentially abundant between the two environments (lymph node vs. primary tumor; q ≤ 0.05; Benjamini-Hochberg test or proteins detected exclusively in one group) **(Table S2-9; Table S3-7)**. Indeed, there is a more heterogeneous proteome composition for non-malignant samples compared to that of the profile detected for the malignant portion of HNSCC, and this may reflect the distinct constitutions of primary tumor and lymph node cells that surround the tumor and metastasis. Differentially abundant proteins are implicated in distinct biological processes compared to those in the malignant samples, and these processes primarily include the upregulation of ribosomal constituents in the lymph node site (Enrichment FDR = 1.63E-37 to 7.36E-30) **(Figure 4B)**. The differential modulation of translation may be related to the heterogeneous cellular composition of tumor and lymph node microenvironments. This constitution reflects the intrinsic function of lymph nodes in the immune response that harbors the transition of naive to effector T lymphocytes and is characterized by the high abundance of the translational machinery and enhanced protein synthesis (Araki et al., 2017; Ricciardi et al., 2018; Xia et al., 2019). As previously reported, the higher translational levels may also be related to the significant enrichment of B cells, T cells, macrophages, neutrophils, plasma cells, myofibroblasts, and specific CAF populations in breast cancer and HNSCC pN+ lymph nodes compared to that at primary sites (Lee et al., 2021; Puram et al., 2017), or these levels may even be associated with the recruitment of neutrophils, monocytes, and macrophages that are necessary for the preparation of pre-metastatic niches in pN0 sites of metastasis (Costa-Silva et al., 2015; Qian et al., 2011; Wculek and Malanchi, 2015).

Next, to visualize the relationship between the cell populations from tumor and nodal sites within the patients in the two populations of malignant and non-malignant cells, we performed a hierarchical clustering analysis. Interestingly, the proteomic profile grouped malignant cells from the two regions by patient, while non-malignant cells were clustered according to site **(Figure 4C-D).** Non-malignant cells derived from sample 400, a tonsil tumor, were clearly grouped with non-malignant samples from the lymph nodes **(Figure 4D)** and exhibited a similar proteomic pattern that has been observed to result from the enhanced lymphoid composition of tonsillar HNSCC.

These analyses clarified the proteome composition that is associated with locoregional metastasis in HNSCC, thus providing an unprecedented view of potential processes and molecules that are implicated in tumor invasion and spread. Again, the results achieved using proteomics resemble the data obtained using scRNASeq for HNSCC (Puram et al., 2017), thus indicating that the methods applied in this study effectively identified and quantified protein content patterns that can be used to reveal the heterogeneity of the populations in the nodal and primary sites.

### Microenvironment proteomes group samples according to metastasis and highlight candidate markers of locoregional spread

We further investigated the potential of multisite proteomic profiles in regard to grouping patients according to lymph node metastasis. Hence, we evaluated the ability of HNSCC global proteomes to separate pN+ and pN0 patients by applying unsupervised hierarchical clustering and associating patients’ clusters (C) with lymph node metastasis (2,451; 1,984; 2,137; 2,188; and 1,154 proteins, respectively, identified across 25 samples of malignant cells from the primary tumor, 27 samples of non-malignant cells adjacent to the primary tumor and lymph node, and 24 buffy coat and saliva samples) **(Figure 5A; Table S2-1 to 7)**. The cluster behavior provides novel insights into the analysis of clinical data and, interestingly, C1 and C2 from lymph node microenvironment cells were non-randomly associated with pN status (p = 0.046; Fisher’s exact test) **(Figure 5B)**. Moreover, clusters were significantly associated with smoking habits (primary tumor and lymph nodes: non-malignant, buffy coat; p = 0.028, p = 0.012, and p = 0.033, respectively), pT (primary tumor - non-malignant and saliva; p = 0.021 and p = 0.050, respectively), desmoplasia status (buffy coat; p = 0.044), and overall survival (saliva cells; p = 0.013) (Fisher’s exact test or log-rank tests) **(Figure S3F-G)**. Based on the knowledge that HPV plays a role in the etiology of HNSCC (primarily in oropharynx squamous cell carcinomas) (Isayeva et al., 2012; Leemans et al., 2018), we also evaluated the presence of this virus in the 27 HNSCC primary tumors to determine its association with patient clustering. HPV16 DNA positivity was detected in 29.6% of HNSCC, including five larynx, two oral, and one oropharyngeal tumor **(Figure S3H)**. Within this subgroup, three tumors from the oropharynx (n = 1), oral (n = 1), and larynx (n = 1) sites were positive for E6/E7 viral transcripts, thus indicating infection by transcriptionally active HPV **(Figure S3H)**. HPV DNA or RNA positivity was not associated with patient clustering patterns (p > 0.05; Fisher’s exact test) (data not shown), thus revealing that other etiological factors were associated with the observed proteomic profiles.

**Figure 5.**
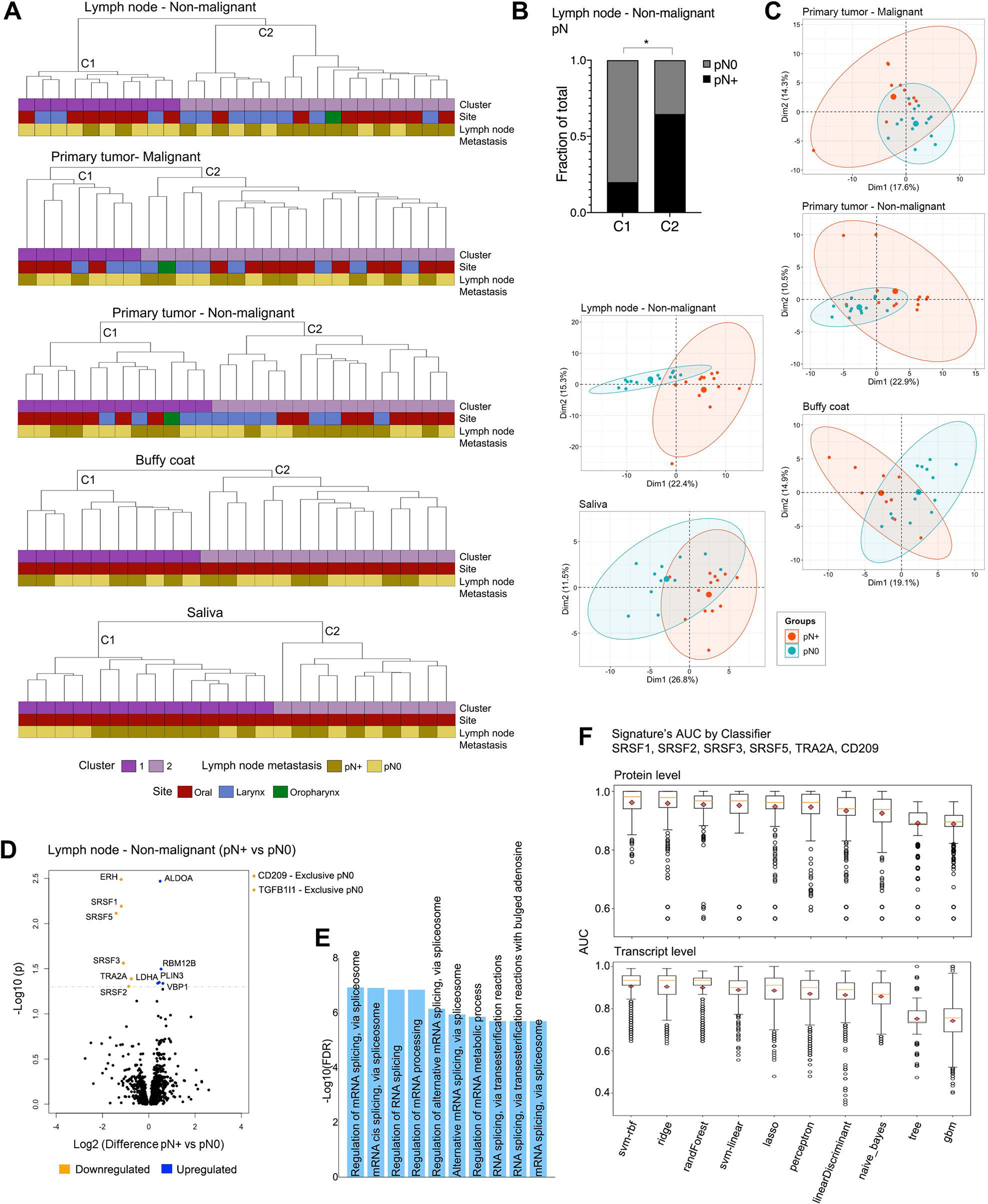
Proteome-grouping pattern associated with nodal metastasis and refinement of targets for ML analysis. (A) Clustering of tissues and fluids from 59 patients based on the global proteomic profile (C1 and C2 clusters). C1 and C2 groups were generated using Complete Canberra (25 primary tumor – malignant samples: 2,451 proteins), Ward Chebyshev (27 primary tumor – non-malignant: 1,984 proteins, 27 lymph node – non-malignant: 2,137 proteins, 24 buffy coat: 2,188 proteins), and Complete Sqeuclidean (24 saliva samples: 1,154 proteins). (B) Association between lymph node metastasis and patient clustering for non-malignant cells from the lymph node (p ≤ 0.05; Fisher’s exact test). *p ≤ 0.05. (C) PCA plots presenting clusters of samples based on the abundance of proteins that were differentially abundant between pN+ and pN0 (p ≤ 0.05; Student’s t-test; 201, 110, 85, 80, 54 differentially abundant proteins from non-malignant cells from lymph nodes, malignant cells from primary tumor, non-malignant cells from primary tumor, buffy coat samples, and saliva samples, respectively). (D) Volcano plot for the differential protein abundance between pN+ and pN0 non-malignant cells from lymph nodes. Differentially abundant proteins are presented as blue and orange dots (pN+ vs pN0; q ≤ 0.05; Benjamini-Hochberg test or proteins detected exclusively in one group in at least 50% of samples). (E) Top-10 GO biological processes that were significantly enriched for the 13 proteins associated with lymph node metastasis from (D) (Enrichment FDR ≤ 0.05). (F) AUC distribution per classifier using ML analysis of SRSF1, SRSF2, SRSF3, SRSF5, TRA2A, and CD209 proteins (upper panel) and transcripts (lower panel). Empty circles indicate data points that were located outside of the whiskers of the box plot (outliers). See also Figures S3-S4.

To further explore the proteomic differences between pN+ and pN0, we employed principal component analysis (PCA) using the sets of deregulated proteins from multiple sites (pN+ vs. pN0; p ≤ 0.05; Student’s t-test or proteins detected exclusively in one group; 201, 110, 85, 80, and 54 differentially abundant proteins from non-malignant cells from lymph nodes, malignant cells from primary tumor, non-malignant cells from primary tumor, buffy coat, and saliva, respectively) **(Figure 2A-B; Table S3-1 to 5)**. The best segregation was observed for the differentially abundant proteins determined for non-malignant cells from lymph nodes that separated the groups according to the first and second components with 22.4% and 15.3% of the total data variation, respectively **(Figure 5C)**.

Based on the observation that non-malignant cells from lymph nodes exhibited the most promising association with lymph node metastasis status according to hierarchical clustering and PCA analysis, we applied FDR to control for multiple hypotheses and to filter out proteins that were strongly associated with the metastatic phenotype (pN+ vs. pN0; q ≤ 0.05; Benjamini-Hochberg test or proteins detected exclusively in one group). Eight downregulated and five upregulated proteins (pN+ vs pN0) had a highly significant differential abundance according to the status of locoregional metastasis (q = 3.25E-03 to 4.91E-02 for nine proteins; two proteins were exclusively detected in 50% of pN0 samples: CD209 and TGFB1I1) **(Figure 5D; Table S3-8)**. This set of 13 proteins is primarily involved in mRNA splicing processes (enrichment FDR = 1.2E-07 to 2.0E-06) **(Figure 5E)**. FDR correction was also applied to define proteins associated with locoregional metastasis in the other four datasets; however, no significant results were observed. We then performed an additional analysis to refine the 13 targets. Downregulation of this set of proteins was associated with other poor prognostic features (p ≤ 0.05; Student’s t-test, ANOVA, and Fisher’s exact test) **(Figure S4A)**, and they were demonstrated to be physically or functionally associated according to protein-protein interaction networks **(Figure S4B).**

Finally, using an ML model, we predicted the power of the five splicing proteins to distinguish pN+ and pN0 patients. We also included the protein CD209 in ML analysis due to the involvement of immune processes in HNSCC that are described at several sites (please see **STAR Methods**). Distinct pairs <Signature Si, Classifier Cj> could discriminate pN+ and pN0 HNSCC patients with elevated AUCs (mean AUC = 0.933), thus indicating the high performance of the six proteins in distinguishing patients according to pN **(Figure 5F; Table S4-1)**. Similarly, transcripts of the six genes also exhibited lower expression in pN+ compared to that in pN0 (RT-qPCR; p = 1.14E-04 to 2.45E-02; Student’s t-test), and this corroborates the proteomic pattern **(Figure S4C-D).** Additionally, transcript pairs <Cj, Si> could discriminate pN+ and pN0 HNSCC patients with high AUCs (mean AUC = 0.853) **(Figure 5F; Table S4-2)**. In summary, we demonstrated the importance of the six metastasis markers in HNSCC clinics and then investigated their relationship with the immune compartment.

### Microenvironment markers may be expressed by immune populations

Based on our data and the knowledge that the lymph node metastasis proteome is deeply associated with the immune response **(Figure 2C-E, Figure 3, Figure S3D),** we next investigated if the six selected targets (SRSF1, SRSF2, SRSF3, SRSF5, TRA2A, and CD209) from non-malignant cells from lymph nodes are likely to be expressed by the immune cell types from the lymph node microenvironments defined in **Figure 3A**. Additionally, we evaluated the gene expression in non-immune populations. From the publicly available scRNASeq data analysis, we observed that the six genes were not cluster markers of immune cells; however, we did determine that they are expressed by the immune milieu from tumors and lymph nodes in an immune cell-type-specific manner **(Figure 6A-C)**. Globally, splicing genes were highly expressed in CD8+ T effector cells and regulatory T cells (Tregs) from pN+ microenvironments (non-malignant cells of tumor and lymph nodes), while *CD209* exhibited the most elevated transcript levels in macrophages **(Figure 6B-C)**. The Treg population exerts a potent immunosuppressive function that plays a crucial role in tumor immune escape (Sakaguchi, 2000). Thus, although we observed a low abundance of the metastasis-dependent splicing proteins in the pN+ environment when compared to pN0, a reducionist view of single cells revealed high expression of this set of genes in Tregs from pN+ primary tumor and lymph node sites, suggesting a role for these molecules in the suppressive environment associated with the metastatic phenotype. We observed a more heterogeneous distribution of immune cell-specific gene expression in the pN0 tumor microenvironment, where *SRSF1* transcripts were primarily depleted in immune cells while other splicing genes (*SRSF2, SRSF3, SRSF5, TRA2A)* were strongly expressed in CD8+ T effector and B cells **(Figure 6A)**. *CD209* exhibited high transcript levels in macrophages **(Figure 6A)**.

**Figure 6.**
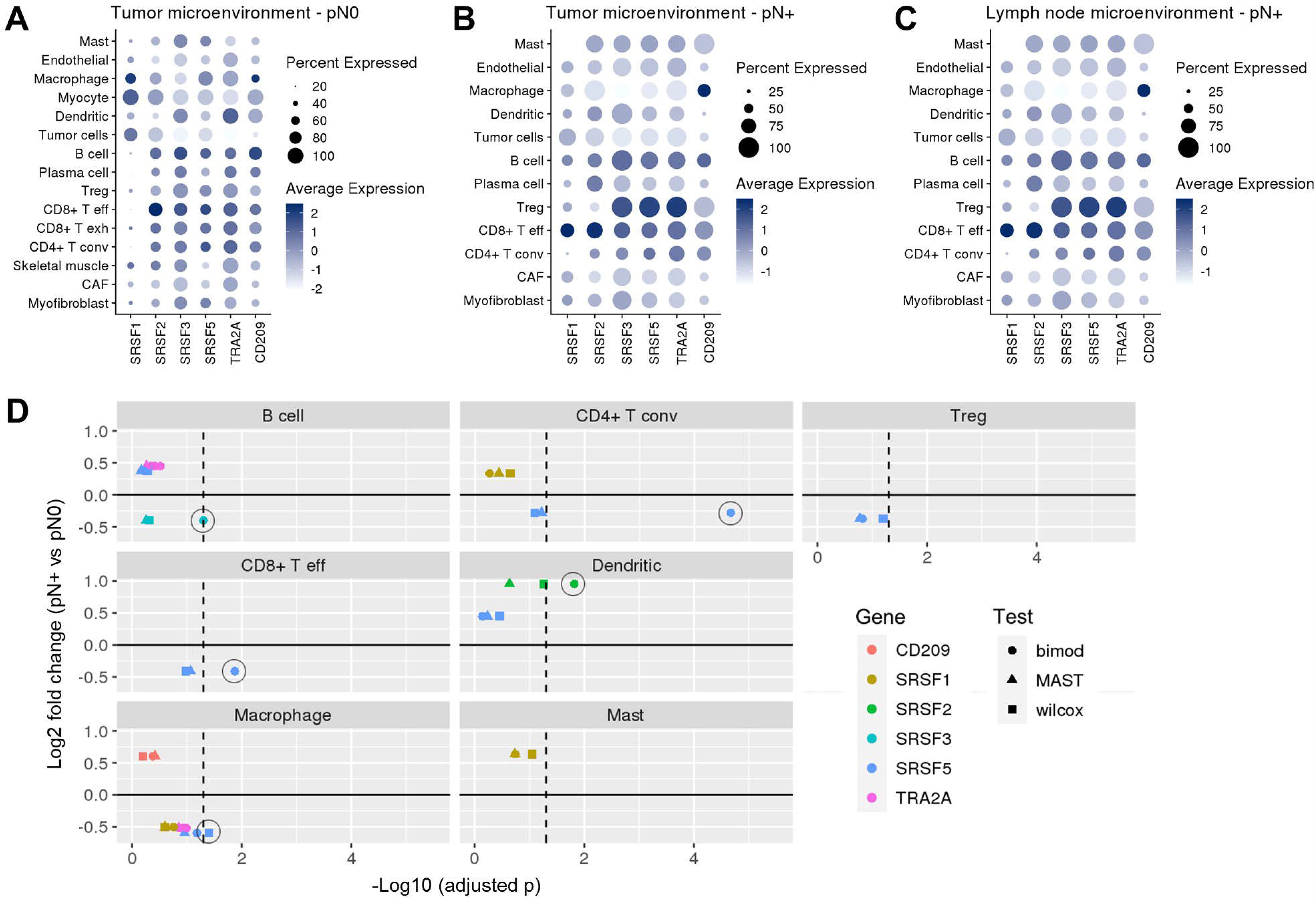
Immune cell-type-specific gene expression associated with six metastasis-dependent markers according to scRNASeq data. (A-C) Expression of the six selected targets (SRSF1, SRSF2, SRSF3, SRSF5, TRA2A, and CD209) in immune populations from HNSCC that were identified by scRNASeq public data (Puram et al., 2017) in primary tumor (A, B) and lymph node (C) microenvironments. (D) Average fold change (y axis) and differential gene expression adjusted p (x axis) between pN+ and pN0 tumor microenvironments for the six targets considering individual immune populations. The analysis was performed with transcript information shown in (A-B) (Puram et al., 2017) and three tests were used to compare the genes for differential expression between pN+ and pN0, including Bimodal (circle), MAST (triangle), and Wilcox (square). The dashed lines represent a p threshold of 0.05 and significant results are labeled with empty circles. See also Figure S3.

When comparing pN+ versus pN0 tumor environments, downregulation of *SRSF3* and *SRSF5* in B cells, CD8+ effector T cells, CD4+ conventional T cells or macrophages was detected **(Figure 6D)**, and the reduced levels in pN+ are in accordance with the proteomic analysis of these sites. Moreover, none of the six microenvironment targets were expressed in CD8+ exhausted T cells from the pN+ tumor microenvironment **(Figure 6D)** even though this cell population has been detected in tumor microenvironments independently of lymph node commitment **(data not shown)**. These results suggest that these immune subpopulations are strong candidates for expressing metastasis-associated candidate markers (p ≤ 0.05; Bimodal, MAST, or Wilcox tests). Moreover, particularly, *TRA2A* was downregulated in the non-immune population of CAFs (pN+ vs. pN0; p ≤ 0.05; Bimodal, MAST, or Wilcox tests) **(Figure S3E)**. Thus, the six selected metastasis-dependent markers were all associated with clinics and presented a value in the immune response, and collectively with the nine immune markers that were previously prioritized, they were evaluated for their ability to discriminate pN+ and pN0 patients using targeted methodologies and an ML model.

### Metastasis-associated tissue targets can be detected in liquid biopsies

Next, we examined if the selected immune (CD3, CD4, CD8, CD11b, CD14, CD16, CD19, CD45, and CD66b) and microenvironment (SRSF1, SRSF2, SRSF3, SRSF5, TRA2A, and CD209) candidate markers could be identified in liquid biopsies using targeted approaches. Liquid biopsies exhibit great potential for cancer management, as they all for the assessment of markers in a minimally invasive manner (Amelio et al., 2020). Moreover, lymph node cells can broadly circulate throughout the body (Blum and Pabst, 2007), and nodal markers can be identified in body fluids as a more convenient source of biomarkers. We used selected reaction monitoring-mass spectrometry (SRM-MS) to measure the relative abundance of selected peptides and proteins in 19 buffy coat (7 pN+, 12 pN0) and 25 saliva samples (16 pN+, 9 pN0) from a 36-HNSCC patient cohort **(Table S5)**. Quality control revealed an average carryover of iRT of 0.16% (0.1%–0.45%) between SRM-MS assays, and this is similar to the 0.1% reported in the literature (Abbatiello et al., 2013) **(Figure S5A)**. Quality parameters **(STAR Methods)** were satisfactory for all evaluated samples **(Figure S5B-E; Figure S6)**. Selected peptides from the 15 immune and microenvironment candidate markers were detected in the buffy coat HNSCC cohort, whereas for saliva samples only the peptides SRSF3_Pep1, SRSF5_Pep1, SRSF5_Pep2, CD45_Pep1, and CD4_Pep1 could not be confidently measured due to low signal-to-noise ratios **(Table S5-2 to 5)**. We also evaluated the gene expression of the six microenvironment targets in a 24-patient cohort from buffy coat samples (10 pN+, 14 pN0) and a 22-patient cohort from saliva cells (10 pN+, 12 pN0) using RT-qPCR. All targets were identified in both fluids **(Tables S5-4 and 5)**. Thus, we demonstrated that blood and saliva contain the metastasis markers that were previously detected for lymph nodes, and we next employed a multiparametric ML approach.

### Machine learning predicts liquid biopsy-metastasis signatures with high performance

Finally, we used an ML strategy to evaluate the power of candidate prognostic markers in predicting nodal status in liquid biopsies. Individual (peptide, protein, or transcript) or combined (peptides + proteins + transcripts) datasets from the 15 immune and microenvironment markers (CD3, CD4, CD8, CD11b, CD14, CD16, CD19, CD45, CD66b, SRSF1, SRSF2, SRSF3, SRSF5, TRA2A, and CD209) assessed in saliva and buffy coat by SRM-MS or RT-qPCR were used for ML analysis **(Figure 7A; Table S4-3 to 15)**. High-performance signatures were defined as stated in the **STAR Methods**. When analyzing individual datasets, we filtered out 24 signature pairs <Si, Cj> from the buffy coat that exhibited a high performance in discriminating pN+ and pN0 patients **(Table S4-3 to 7 and 13)**. SRSF1 protein was the most frequent target among the high-performance signatures within <Si, Cj> (n = 10 signatures) **(Figure 7B)**. Proteins CD45 and SRSF1 were defined as high-performance signatures according to the largest number of classifiers (CD45: lasso, linear discriminant, ridge, and linear SVM; SRSF1: lasso, perceptron, ridge, and linear SVM) **(Figure 7C)**. The top-1 pair <Si, Cj> (highest average AUC) that most effectively differentiated pN+ and pN0 patients among buffy coat individual datasets was based on the combined abundance of three immune peptides using a random forest classifier: CD8_Pep1, CD11b_Pep2, CD11b_Pep3 (AUC = 0.933; 95% CI = 0.91 – 0.96; sensitivity = 0.783; specificity = 0.933) **(Figure 7D; Figure S7A)**. Eighteen high-performance signatures for <Si, Cj> were observed in saliva cells with a predominance of CD11b_Pep1, CD16_Pep1, CD45_Pep2, and TRA2A_Pep1 (5 signatures/each) **(Table S4-8 to 13; Figure 7B)**. Protein CD14 was observed to be a high-performance signature according to the largest number of classifiers (linear discriminant, ridge, linear SVM, and RBF SVM) **(Figure 7C)**. TRA2A protein abundance using random forest classifier could discriminate patients based on nodal status with the highest AUC (AUC = 0.868; 95% CI = 0.86 – 0.88; sensitivity = 0.869; specificity = 0.811) **(Figure 7D; Figure S7B)**.

**Figure 7.**
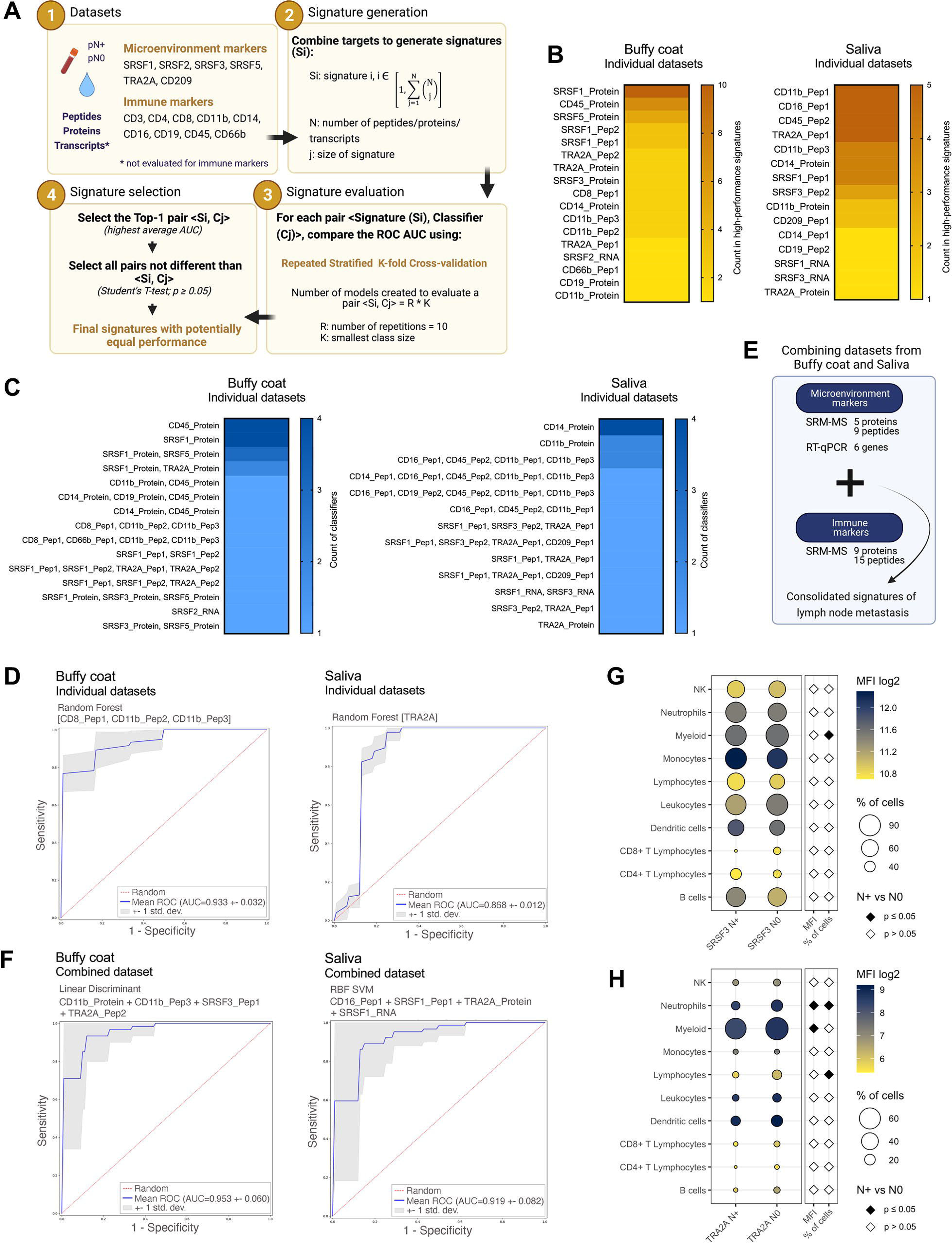
Definition of the best prognostic signatures in liquid biopsies according to machine learning. (A) Design used to determine the signatures of lymph node metastasis with the best performance according to ML. AUC: area under the curve. (B) Count of microenvironment and immune markers in buffy coat (left panel) and saliva (right panel) signatures that exhibited high performance in regard to discriminating pN+ and pN0 patients. Individual peptide, protein, or transcript datasets were considered. (C) Count of signatures per classifier in buffy coat (left panel) and saliva (right panel) samples. Only signatures exhibiting high performance for discriminating pN+ and pN0 HNSCC patients were included. Individual peptide, protein, or transcript datasets were considered. (D) ROC curves indicating the top-1 pair <Si, Cj> possessing higher AUC across all datasets evaluated in buffy coat (left panel) and saliva (right panel) samples. Individual peptide, protein, or transcript datasets were considered. AUC: area under the curve. (E) Experimental approach to define the best consolidated lymph node metastasis signatures using ML. (F) ROC curves indicating the top-1 pairs <Si, Cj> in buffy coat (left panel) and saliva (right panel) samples that discriminated HNSCC patients based on lymph node metastasis status (AUC > 0.919). Combined microenvironment and immune peptide, protein, and transcript datasets were considered. (G-H) Log_2_ mean fluorescence intensity (MFI) and percentage of immune cells expressing the proteins SRSF3 (G) and TRA2A (H) in N+ and N0 buffy coat samples (10 N+; 10 N0). Significant differences between groups are represented as black diamonds (Student’s t-test, Welch’s t-test or Mann-Whitney test; p ≤ 0.05) See also Figures S5-S7.

We then combined all single molecular datasets to generate a consolidated signature of the tumor microenvironment and immune markers using peptide, protein, and transcript (when available) information from buffy coat or saliva **(Figure 7E; Table S4-13 to 15)**. These analyses yielded higher performance in regard to separating pN+ and pN0 compared to that of the individual datasets as evidenced by AUCs > 0.9 **(Figure 7F; Table S4-13)**. Top-1 signatures could discriminate pN+ and pN0 patients with elevated AUCs (AUC > 0.92) and included four targets for buffy coat (CD11b_Protein, CD11b_Pep3, SRSF3_Pep1, and TRA2A_Pep2; linear discriminant; AUC = 0.953; 95% CI = 0.91 – 1.00; sensitivity = 0.933; specificity = 0.907) and four targets for saliva (CD16_Pep1, SRSF1_Pep1, TRA2A, *SRSF1* transcript; RBF SVM; AUC = 0.919; 95% CI = 0.86 – 0.92; sensitivity = 0.834; specificity = 0.936) **(Figure F; Figure S7C-D)**. Based on this analysis, we defined sets of targets for combined peptide/protein/transcript datasets that significantly outperformed lymph node metastasis markers identified using single datasets.

Finally, to indicate the specific circulating immune populations responsible for the top-1 combined signature detected in buffy coat (SRSF3 and TRA2A proteins), we performed flow cytometry analysis in blood samples from an independent 20-HNSCC patient cohort (10 N+; 10 N0) **(Figure S7E-F)**. Both proteins were highly expressed in an elevated proportion of buffy coat’s myeloid immune populations **(Figure 7G-H)**. N+ samples had a lower proportion of myeloid cells expressing SRSF3 when compared to N0 (p < 0.001; Mann-Whitney test) **(Figure 7G)**, as well as downregulation or a reduced proportion of myeloid cells (p = 0.050; Student’s t-test), neutrophils (p = 0.046; Student’s t-test and p = 0.050; Student’s t-test), and lymphocytes (p = 0.043; Welch’s t-test) expressing TRA2A **(Figure 7H)**. Notably, the reduced levels of SRSF3 and TRA2A in the circulating cells of N+ patients reflect the lower abundance of these proteins found in the pN+ microenvironment of lymph nodes (**Figure 5D**). Moreover, the relevance of neutrophils detected for TRA2A is in line with the enrichment of neutrophils-dependent biological processes identified for metastasis markers in buffy coat **(Figure 2C),** strengthening the significance of this immune population in the metastatic cascade.

Taken together, the results of this study depicted the framework of the wired microenvironments in HNSCC and provide a promising basis for understanding tumor biology, indicating a potential signature of metastasis for this disease.

## DISCUSSION

Cancer is a systemic disease, and the crucial contribution of multiple environments to the regulation of HNSCC implicates a fundamental role of varied populations in supporting a tumoral niche. Hence, further progress in head and neck oncology will require a global understanding of the diverse molecular landscape of the neoplasm. Additionally, molecular profiling of isolated cell populations from tissues reduces the intrinsic heterogeneity caused by the mixing of cell types and is essential for deciphering cancer. Therefore, we performed a comprehensive mass spectrometry-based proteomic analysis of isolated cell populations from tissues and fluids to characterize HNSCC.

Our key findings are in regard to the modulation of the immune system retrieved from the global and metastasis-dependent HNSCC proteomes **(Figure 1C-E; Figure 2C-E; Figure 3; Figure 6; Figure S3D)**. The tumor immune infiltrating composition has been previously explored in HNSCC (Mandal et al., 2016; Nordfors et al., 2013; Trellakis et al., 2011); however, a breakthrough in our study is the identification of the overrepresentation of immune processes in the multisite samples evaluated. Remarkably, the immune response is coordinated across tissues, and its relationship to cancer must encompass the peripheral immune system in addition to the TME (Hiam-Galvez et al., 2021). The association between the immune population and global proteomes identified here is of special interest, as the immune context exerts profound effects on the response to immunotherapies. Resistance to immunotherapy remains a bottleneck in regard to the successful treatment of cancer, and greater than 80% of HNSCC patients with metastatic disease do not respond to PD-1 blockade (Bauml et al., 2019). Hence, biomarkers for patient stratification or the identification of new immune targets are urgently needed. The multisite proteomic analysis presented in this study elucidated the immune cell composition of HNSCC TME and blood **(Figure 1D-E)**, and this is of paramount relevance to guide further therapeutic strategies. Additionally, our study revealed two biological processes that are highly enriched for clusters of proteins, and these processes include antigen processing and presentation (APP) and the Fc receptor signaling pathway (**Figure 1C)**. Alterations in the APP are known to be associated with immune evasion and can result in impaired antitumor responses and therapy resistance (Gettinger et al., 2017; Sade-Feldman et al., 2017). Additionally, blockade of the interaction between the inhibitory Fc receptor FcγRIIB and immune complexes in mouse models has been proposed as an approach for cancer immunotherapy (Kasahara et al., 2019). Thus, we identified immune cell players and components of specific pathways that can be further targeted to enhance the immunogenicity in HNSCC or that can be further explored as markers of immunotherapy resistance.

Understanding the molecular principles underlying cancer invasion and metastasis is a highly complex endeavor and is of special interest in HNSCC due to its association with poor prognosis (Ho et al., 2017). Remarkably, our work revealed that pN status is clearly related to immunity in HNSCC by using a combination of our own proteome data, public RNASeq information, and cell counts obtained from clinical laboratories **(Figure 2C-E; Figure 3; Figure 6; Figure S3D)**. By deciphering the immune GO biological processes associated with metastasis markers that are commonly enriched in multiple sites, we revealed that the development of lymph node metastasis depends upon an orchestrated interconnection of immune system across multiple HNSCC environments **(Figure 2C-E)**. This interconnection may be necessary to maintain an immunosuppressive systemic environment in HNSCC that is a prerequisite for preparing the pre-metastatic niche and thus allowing for the establishment and maintenance of cancerous cells in the pN+ lymph nodes (Jones et al., 2018). Indeed, a suppressed pN+ environment may be supported by a group of splicing proteins that are strongly associated with the metastatic phenotype in our HNSCC dataset, and the genes associated with these proteins include *SRSF1, SRSF2, SRSF3, SRSF5*, and *TRA2A,* and also the immune marker *CD209* **(Figure 5D).** While the splicing genes are highly expressed in the immunosuppressive Treg cell population of the pN+ tumor and lymph node microenvironments, *CD209* exhibits high transcript levels in pN+ macrophages that can potentially prevent CD8+ T cells from exerting their full cytotoxicity against tumor cells **(Figure 6)** (Cassetta and Kitamura, 2018).

Additionally, lymphatic metastasis is the primary route for the development of distant metastasis in a subset of tumors (Naxerova et al., 2017; Zhang et al., 2020), and blocking the spread of tumor cells via lymphatic vessels may prolong the life of cancer patients and also mitigate the poor prognosis. By comparing malignant and non-malignant cells between primary sites and lymph nodes **(Figure 4)**, we identified several proteins involved in the metastatic spread that can be targeted for therapy, including motility- and translation-associated molecules **(Figure 4A-B)**. Indeed, the hierarchical clustering analysis revealed that non-malignant cells possess a more homogeneous proteomic profile across patients than do malignant cells when the samples are grouped according to site instead of by patient **(Figure 4C-D)**. The same pattern was observed at the scRNA level in HNSCC (Puram et al., 2017), thus indicating that targeting the microenvironment cells may yield improved treatment responses due to its reduced dynamism across patients.

Although a long list of prognostic biomarker candidates can be observed in the literature, no molecular marker has been widely accepted for routine use in managing patients with HNSCC. Currently, only HPV and p16^INK4A^ are considered to be prognostic markers for oropharyngeal cancer, and both are associated with an improved prognosis (National Comprehensive Cancer Network, 2011). Hence, it appears that a multiparametric assessment will be necessary to dissect the complex tumor interactions and define prognosis in HNSCC, and this assessment will primarily include lymph node metastasis-dependent signatures that represent the primary poor prognosis feature. In this scenario, the multisite analysis led us to develop a multiparametric ML model that allowed for the indication of high-performance signatures of peptides, proteins, and/or transcripts in saliva or blood that can discriminate pN+ and pN0 HNSCC (AUC > 0.868 for saliva and AUC > 0.933 for blood) **(Figure 7; Figure S7)**. We successfully combined multiple datasets for ML analysis (peptides + proteins + transcripts for immune and microenvironment markers), and the top-1 signatures for buffy coat samples (CD11b_Protein, CD11b_Pep3, SRSF3_Pep1, and TRA2A_Pep2; linear discriminant; AUC = 0.953) and saliva samples (CD16_Pep1, SRSF1_Pep1, TRA2A, *SRSF1* transcript; RBF SVM; AUC = 0.919) clearly outperformed signatures indicated by individual datasets in regard to separating patients according to pN status. Interestingly, evaluating the expression of the top-1 signature defined for buffy coat (SRSF3 and TRA2A proteins) in cells from blood **(Figure 7G-H)** strengthened the paramountcy of the myeloid lineage, especially neutrophils, in the metastatic phenotype of HNSCC **(Figure 2C)**. Nevertheless, herein we show a disease snapshot and whether the modulation of neutrophils is a cause or effect of HNSCC dissemination needs further investigation. In summary, these metastasis-dependent signatures are promising once they can non-invasively guide decision making for HNSCC patients, and further verification in larger cohorts may confirm their clinical utility.

In conclusion, our results explored the molecular mechanisms underlying HNSCC carcinogenesis through a multisite quantitative mass spectrometry-based proteomics analysis that identified potential cell players and biological processes largely related to immune system that can be further targeted for therapy or explored as prognosis markers. High-performance locoregional metastasis-dependent signatures have been depicted and are promising for future clinical implementation. Herein, we investigated the HNSCC multisite landscape in an unprecedented manner, and our results provide the clinical and research communities with valuable information that may guide the management of patients.

## ACKNOWLEDGMENTS

This work was supported by FAPESP under Grant numbers 2010/19278-0, 2016/07846-0, and 2018/18496-6 to A.F.P.L., 2015/19191-6 and 2019/21815-9 to A.F.B.L., CNPq Grant number 305851/2017-9 to A.F.P.L, and ANID-FONDECYT Grant number 1190775 to W.G.A. This research used the proteomics facility of the Brazilian Biosciences National Laboratory (LNBio) that is part of the Brazilian Center for Research in Energy and Materials (CNPEM), a private non-profit organization under the supervision of the Brazilian Ministry for Science, Technology, and Innovation (MCTI). The mass spectrometry laboratory staff are acknowledged for their assistance during the experiments (Proposal number MAS-22044). We also acknowledge Prof. Dr. Tsai Siu Mui, CENA, USP, for the use of the Leica Laser Microdissection System LMD6 (FAPESP Grant number 2009/53998-3), Waters Corporation for providing access to the Acquity UPLC-Class M system coupled with a Xevo TQ-XS triple quadrupole mass spectrometer, Prof. Guilherme Telles, IC, UNICAMP, for his support with the selection of proteotypic peptides for SRM-MS analysis, and the Laboratory for Integrative and System Biology (LaBIS) for the use of the LaBIS Cloud (FAPESP Grant numbers 2011/00417-3 and 2015/50612-8). The Graphical Abstract and Figures 7A and 7E were created with BioRender.com.

## AUTHOR CONTRIBUTIONS

Conceptualization, A.F.B.L., A.F.P.L.; Methodology, A.F.B.L., A.F.P.L., Formal Analysis, A.F.B.L., F.M.S.P., H.H., M.A.P.; Investigation, A.F.B.L., C.R., L.X.N., D.C.G., T.R.M., A.G.C.N., R.R.D., B.P.M., P.A.L., N.A.L.G.; Resources, A.N.R., N.K.C., K.J.G., L.L.V., M.U., A.C.P.R, T.B.B., M.B., W.G.A., A.F.P.L.; Writing – Original Draft, A.F.B.L.; Writing – Review & Editing, A.F.B.L., A.F.P.L., T.S.M, L.L.V., L.P.K.; Supervision, A.F.P.L.; Funding Acquisition, A.F.P.L.

## DECLARATION OF INTERESTS

The authors declare no competing interests.

## STAR METHODS

### RESOURCE AVAILABILITY

#### Lead Contact

Further information and requests for resources and reagents should be directed to and will be fulfilled by the lead contact, Adriana Paes Leme.

#### Materials availability

This study did not generate new unique reagents.

#### Data and code availability

The mass spectrometry DDA proteomics data generated in this study are available at ProteomeXchange via the PRIDE partner repository with the dataset identifier PXD027780 (http://www.proteomexchange.org/) (Perez-Riverol et al., 2019). SRM-MS data for the measured proteins are available through the Panorama repository at the link https://panoramaweb.org/16fpvB.url and ProteomeXchange dataset identifier PXD027984.

### EXPERIMENTAL MODELS AND SUBJECT DETAILS

#### Patients and sampling

An 83-patient cohort with HNSCC obtained from the oral cavity (n = 71 patients), larynx (n = 11 patients), and oropharynx (n = 1 patient) was included in this study **(Table S1)**. Informed consent was obtained from all individuals as approved by the Ethics Committees of Carlos Van Buren Hospital (Process 121), University of Valparaíso (Process CB051-14), University of São Paulo Academic Biobank of Research on Cancer, Centro de Investigação Translacional em Oncologia, Instituto do Câncer do Estado de São Paulo (ICESP) (Protocol CAEE 30658014.1.1001.0065), Faculty of Medicine of Jundiai (Protocol CAEE 45091715.1.0000.5412), and A.C. Camargo Hospital (Protocol 2532/18B). The 83 patients were separated into two groups that included discovery and verification cohorts. The discovery cohort consisted of 59 HNSCC patients from whom 27 FFPE primary tumors, 27 FFPE lymph nodes, 24 blood, and 24 saliva samples were collected and used in discovery proteomics (DDA) **(Table S1)**. This group included 27 matched tumors and lymph node tissues from 27 HNSCC individuals and also 15 paired buffy coats and saliva samples recovered from 32 patients. The verification cohort comprised a 79-patient group, and of these, 19 FFPE lymph nodes, 47 blood, and 33 saliva samples were used in the SRM-MS, RT-qPCR, and flow cytometry experiments. Of the 79 patients used in the verification, 34 were independent cohorts that were not evaluated in discovery proteomics. Saliva was collected as previously described (Carnielli et al., 2018), and collection occurred preferably in the morning from individuals who had not eaten or ingested liquids (except water) and had undergone oral hygiene at least 1 h prior to collection. Patients were instructed to rinse their mouths with 5 mL of drinking water, and saliva was subsequently harvested without stimulation into a plastic receptacle. Four mL of peripheral blood were collected in BD Vacutainer® tubes with EDTA as an anticoagulant (BD Life Sciences, USA). Detailed sample sizes used in each methodology are presented in **Table S1-1**, listed below, or presented throughout the text. The cases were histopathologically classified according to the recommendations of the World Health Organization (Barnes et al., 2005), the American Joint Committee on Cancer (AJCC), and the International Union Against Cancer (IUAC) (Amin et al., 2017). None of the patients received neoadjuvant radiotherapy or chemotherapy prior to sample collection. The clinical and pathological data are summarized in **Table S1-2**.

### METHOD DETAILS

#### Isolation of cells from tissues

Malignant and non-malignant cells were harvested from FFPE primary tumor and lymph node samples using micro- or macrodissection. Histology-guided laser microdissection was used to recover malignant and/or adjacent non-malignant portions from FFPE primary tumors (n = 27 samples) and metastatic lymph nodes (malignant cells; n = 13 samples). Malignant cells from tumors were retrieved from the invasive front, an area with high prognostic potential where the most invasive and aggressive cells reside (Bryne, 1998). The invasive front contains the most advanced tumor cells that invade normal tissues such as muscle, connective tissue, salivary glands, and blood vessels (Bryne, 1998). As described previously (Carnielli et al., 2018), the tumor invasive front was collected from the farthest area of the invasive surface of the tumor, and it was collected up to a depth of one mm in the histological section. Hematoxylin and eosin-stained histological slides were cut into 5 sections to guide microdissection. Slices (10 μm) were mounted on Arcturus PEN Membrane Glass Slides (Life Technologies, USA), deparaffinized with xylene, hydrated in a graded ethanol sequence, and stained with hematoxylin for 1 min. An area of approximately 3,000,000 μ was dissected from one to three slides per sample for (i) malignant cells from primary tumors, (ii) non-malignant cells adjacent to primary tumors (mucosal margins), (iii) malignant cells from lymph nodes, and (iv) non-malignant cells from metastatic lymph nodes adjacent to the metastasis using Leica LMD6 equipment (Leica Biosystems, Germany). Samples were collected in 600 µL tubes. Non-malignant cells from non-metastatic lymph nodes (n = 14 samples) were recovered from 10 μm slices mounted on standard slides, and they were used as the unique site for macrodissection due to the population homogeneity of the tissue. A 3,000,000 μm area was selected according to a comparison to parallel hematoxylin and eosin stained 5 μm sections, scraped into 600 µL tubes, deparaffinized with xylene, and hydrated in a graded ethanol sequence. All dissected samples were stored at − −80 °C until MS analysis.

#### Isolation of cells from fluids

The buffy coat and saliva fractions were obtained using centrifugation. Saliva samples (n = 38 samples) from HNSCC patients were centrifuged for 5□min at 1,500□ × g and 4°C to separate the cells. Peripheral blood samples (n = 52) were centrifuged for 8 min at 1,000□ × g at room temperature, and the cellular portion (buffy coat) was washed with lysis buffer to eliminate red cells (10 mM Tris-HCl pH 7,6, 5 mM MgCl_2_, 10 mM NaCl). For flow cytometry analysis, buffy coat samples were fixed with 4% paraformaldehyde prior to red cell lysis. The cellular portions from the fluids were frozen at −80 °C for further analysis. The buffy coat is primarily composed of leukocytes and granulocytes (Sutton et al., 1988), while saliva samples possess high levels of epithelial cells, leukocytes, and microorganisms (Dewhirst et al., 2010; Theda et al., 2018).

#### Sample preparation for discovery proteomics

Malignant cells from tumors (n = 27 samples), non-malignant cells from tumors (n = 27 samples), malignant cells from lymph nodes (n = 13 samples), non-malignant cells from lymph nodes (n = 27 samples), buffy coat samples (n = 24 samples), and saliva cells (n = 24 samples) were submitted to discovery proteomics. Proteins were isolated from buffy coat samples and saliva cells using TRIzol reagent (Invitrogen, USA) and resuspended in 200□μL of urea buffer (100□ M Tris-HCl pH 7.5, 8□M urea, and 2□M thiourea). Ultrasound treatment was performed for 10 min in an iced water bath for homogenization. Micro- or macrodissected tissue samples were transferred to an 8 M urea solution. All tissue and fluid samples were treated with 5 mM dithiothreitol for 25 min at 56°C and 14 mM iodoacetamide for 30 min at room temperature for cysteine reduction and alkylation, respectively. Urea was diluted to a final concentration of 1.6 M with 50 mM ammonium bicarbonate, and 1 mM calcium chloride was added to the samples. For protein digestion, a total of 2.5 μg of trypsin (Promega, USA) was added in three steps that included 1 μ for 16h, 1 μg for an additional 16h, and 0.5 μg for the last 16h at 37°C. The reactions were quenched with 0.4% formic acid, and the peptides were desalted with C18 stage tips (3M, USA) (Rappsilber et al., 2007) and dried in a speed-vac instrument. Tissue samples were reconstituted in 0.1% formic acid that was applied proportionally to the dissected area (10 μL of formic acid for 1,000,000 μm^2^), and iRT peptides were added to the digested tissue sample at a final concentration of 11.1 fmol/μL for LC-MS/MS quality control (Pierce™ Peptide Retention Time Calibration Mixture, Thermo Scientific, USA). A volume of 4.5 μL was used (50 fmol of iRT peptides). Peptides from the buffy coat and saliva samples were quantified using the Pierce™ Quantitative Colorimetric Peptide Assay (Thermo Scientific, USA), and 2 μg of these proteins were subjected to LC-MS/MS analysis.

#### Discovery proteomics and data analysis

Tissue and fluid samples were analyzed by LC-MS/MS using an ETD-enabled Orbitrap Velos mass spectrometer (Thermo Fisher Scientific, USA) connected to an EASY-nLC system (Proxeon Biosystem, USA) through a Proxeon nanoelectrospray ion source. Peptides were subsequently separated in a 2%–90% acetonitrile gradient in 0.1% formic acid using a PicoFrit analytical column (20□cm ×□ID75, 5□µm particle size, New Objective) at a flow rate of 300□nL/min over a 212 min gradient (35% acetonitrile at 175 min) for tissue and buffy coat samples or a 170 min gradient (35% acetonitrile at 123 min) for saliva samples. The nanoelectrospray voltage was set to 2.2 kV, and the source temperature was 275°C. All instrument methods were configured for data-dependent acquisition (DDA) in the positive ion mode. Full scan MS spectra (*m*/*z* 300-1,600) were acquired in the Orbitrap analyzer after accumulation to a target value of 1e6 ions. Resolution in the Orbitrap was set to r=60,000, and the 20 most intense peptide ions with charge states ≥ 2 were sequentially isolated with an isolation window of *m/z* 3 to a target value of 5,000 and then fragmented in the linear ion trap by low-energy CID (normalized collision energy of 35%). The signal threshold for triggering an MS/MS event was set at 1,000 counts. Dynamic exclusion was enabled with an exclusion size list of 500, an exclusion duration of 60 s, and a repeat count of 1. An activation of q = 0.25 and an activation time of 10 ms were used. Proteins were identified using MaxQuant v.1.5.8.0 (Cox and Mann, 2008; Cox et al., 2011) against the Uniprot Human Protein Database (92,646 protein sequences, 36,874,315 residues, release May 2017) using the Andromeda search engine. Carbamidomethylation was set as fixed modification, and N-terminal acetylation and oxidation of methionine were used as variable modifications. Maximum 2 Trypsin/P missed cleavage, a tolerance of 4.5 ppm for precursor mass, and a tolerance of 0.5 Da for fragment ions were set for peptide identification. Protein groups (also referred to as proteins in the text) were automatically inferred by the Andromeda engine using the parsimony principle. A maximum of 1% FDR calculated using reverse sequences was set for both protein and peptide identification. Protein quantification was performed using the LFQ algorithm implemented in MaxQuant software to reflect a normalized protein quantity deduced from razor+unique peptide intensity values. A minimal ratio count of one and a 2-min window for matching between runs were both required for quantification. Protein identifications assigned as ‘Reverse’ were excluded from further analysis. Contaminants were not removed from the dataset, as keratins are of special interest in the study of squamous epithelial cells. LFQ intensities were log_2_ transformed in Perseus v. 1.3.0.4 (Tyanova et al., 2016) and used in subsequent analyses.

#### Quality control in discovery proteomics

The quality of discovery proteomics assays for tissues and fluids was evaluated by measuring deviations in the retention time for three trypsin autolysis peaks at *m*/*z* 421.7584 +2, 523.2855 +2, and 737.7062 +3. Moreover, four iRT peptides that were spiked into tissue samples were selected for verification of the retention time, intensity normalized by mean per sample, and intensity normalized by mean per group (iRT_Pep1: SSAAPPPPPR; iRT_Pep2: HVLTSIGEK; iRT_Pep3: GISNEGQNASIK; iRT_Pep4: IGDYAGIK) (Pierce™ Peptide Retention Time Calibration Mixture, Thermo Scientific, USA). The peaks were evaluated in Skyline 19.1 (MacLean et al., 2010) using the MSstats tool (Choi et al., 2014). Samples with deviated or recurrent absent intensities/retention times were excluded from the analysis.

#### HPV genotyping

HPV genotyping was performed in 27 primary tumors from HNSCC patients that were included in the discovery proteomics analysis using the INNO-LiPA HPV Genotyping kit in an Autoblot 3000 system (Fujirebio, Japan). HPV positivity was achieved for primary tumors and used in all tissue analyses and once matched tumoral and lymph node samples from the same patients were included in this study. Viral infection was not assessed in HNSCC patients used for buffy coat and saliva analysis due to the unavailability of primary tumor tissues. Sections (10 µm) were cut from FFPE blocks, deparaffinized in xylene, and rehydrated in a graded ethanol sequence. Genomic DNA was extracted using a standard protocol with proteinase K, phenol/chloroform, and ethanol treatment. DNA samples were quantified by spectrophotometry (NanoDrop ND-1000, NanoDrop Technologies, USA). INNO-LiPA is a line probe assay based on the principle of reverse hybridization for the identification of 32 different HPVs, including 13 high-risk genotypes. Biotinylated consensus primers (SPF10) were used to amplify a 65-bp region within the L1 region of multiple HPV types, and the resulting biotinylated amplicons were denatured and hybridized with specific oligonucleotide probes. A primer set for the amplification of human *HLA-DPB1* was used to monitor sample quality and extraction. The tests were performed according to the manufacturer’s instructions.

#### Expression profiling of HPV

HPV DNA-positive tumors (n = 10 samples) were evaluated for the expression of viral transcripts using RT-qPCR. Total RNA was extracted from 10 µm FFPE sections using TRIzol reagent (Invitrogen, USA) and quantified using a Nanodrop ND-1000 spectrophotometer (NanoDrop Technology, USA). RNA was reverse transcribed using the High-Capacity cDNA Reverse Transcription Kit (Thermo Scientific, USA). PCR amplification was performed in an ABI Prism 7500 Sequence Detection System (Applied Biosystems, USA) using SYBR Mix (Applied Biosystems, USA). Two primer pairs were used to amplify E6 gene expression, and two pairs were employed to analyze E7 transcripts from HPV16 (Gao et al., 2013) **(Table S6-1)**. *GAPDH* was used as a control for the sample quality. PCR reactions were performed in duplicate for a total volume of 12.5 μL. Samples with amplification of at least one primer pair for E6 or E7 were considered positive for HPV expression. Two samples could not be evaluated due to poor FFPE RNA quality.

#### Clustering global proteome datasets

The overlay among proteins identified in the proteomes was visualized by upset plots generated using the Intervene tool (Khan and Mathelier, 2017). Hierarchical clusters were generated in Python v3.6 using log_2_ LFQ intensities from the tissues and fluids datasets. Missing values were replaced by random numbers drawn from a normal distribution with a width of 0.3 and a down shift of 1.8. The normal distribution was generated according to negative ranking of mean and standard deviation (μ’, σ’) from columns using the formulas μ’ = μ column – shift × σ column, shift = 1,8; σ’ = width × σ column, width = 0,3. Hierarchical clustering measurements were performed using fastcluster and SciPy 1.0 packages (Müllner, 2013; Virtanen et al., 2020). Methods (’complete’, ‘weighted’, and ‘ward’) and metrics (’braycurtis’, ‘canberra’, ‘chebyshev’, ‘cityblock’, ‘correlation’, ‘cosine’, ‘dice’, ‘euclidean’, ‘hamming’, ‘jaccard’, ‘jensenshannon’, ‘kulsinski’, ‘mahalanobis’, ‘yule’, ‘matching’, ‘minkowski’, ‘rogerstanimoto’, ‘russellrao’, ‘seuclidean’, ‘sokalmichener’, ‘sokalsneath’, and ‘sqeuclidean’) were combined, and dendrograms exhibiting the most evident clustering of proteins (protein clusters PC_n_) or patients (clusters C_n_) were selected for further analysis. Heat maps were drawn using the clustermap tool from the seaborn v.0.11.1 package (Waskom, 2020).

#### Annotation of clusters and association with clinical data

Meaningful GO biological processes that were significantly enriched in protein clusters for blood and tissue datasets were selected using the GSEApy v.0.9.18 package in Python v3.7 against the ‘GO_Biological_Process_2018’ background (adjusted p ≤ 0.05) (Fang, 2020). Fisher’s exact test was employed to associate patient clusters C_n_ with clinical and pathological features in the IBM SPSS Statistics 28.0 software (IBM Corp.), and the association with survival was evaluated using Kaplan-Meier survival curves and the log-rank test (p ≤ 0.05). For fluids, we considered clinical characteristics for comparison that included age, sex, smoking habits, alcohol consumption, vital status (dead or alive), pathologic N (pN) (Amin et al., 2017), pathologic T (pT) (Amin et al., 2017), pathologic stage (Amin et al., 2017), overall survival, lymphatic invasion, perineural invasion, vascular invasion, depth of invasion, desmoplasia, inflammatory infiltrate, and surgical margin status. HPV DNA, HPV RNA, age, sex, anatomical site of the tumor, vital status (dead/alive), smoking habits, alcohol consumption, overall survival, disease-free survival, surgical margin status, recurrence, pT (Amin et al., 2017), pN (Amin et al., 2017), pathologic stage (Amin et al., 2017), depth of invasion, histological grade according to WHO (Barnes et al., 2005), invasion pattern, inflammatory response, degree of keratinization, nuclear pleomorphism, and perineural invasion were used for tissue analysis. Significant associations were visualized using GraphPad Prism v8.2.1 (GraphPad; https://www.graphpad.com).

#### Definition and annotation of metastasis signatures

Log_2_ LFQ intensity values were used to determine differentially abundant proteins between pN+ and pN0 conditions in Perseus v. 1.3.0.4 software (p ≤ 0.05; Student’s t-test and Benjamini-Hochberg test) (Tyanova et al., 2016). Differentially abundant proteins exhibiting higher expression in pN+ compared to that in pN0 or those exclusively detected in pN+ were termed “upregulated”, while proteins exhibiting lower expression in pN+ compared to that in pN0 or those exclusively detected in pN0 were termed “downregulated”. The datasets that were analyzed included primary tumor – malignant cells (11 pN+, 14 pN0 patients), primary tumor and lymph node – non-malignant cells (13 pN+, 14 pN0), buffy coat samples (11 pN+, 13 pN0), and saliva cells (13 pN+, 11 pN0). The intersection among datasets was visualized in upset plots using UpSet (Lex et al., 2014), and STRING was employed to retrieve protein-protein interaction networks (Szklarczyk et al., 2019). The statistical significance and magnitude of changes are represented in volcano plots (R v3.6.2). GO biological processes were enriched using ShinyGO v. 0.61 (FDR ≤ 0.05) (Ge et al., 2020), and the top-10 overrepresented GO terms were simultaneously visualized (Bonnot et al., 2019). Samples were grouped according to the protein profile by applying PCA using the ggfortify v0.4.11 package in R v3.6.2. The relationship between protein abundance and clinicopathological data was determined for the following features: HPV DNA, HPV RNA, age, sex, anatomical site of tumor, vital status (dead/alive), smoking habits, alcohol consumption, overall survival, disease-free survival, surgical margin status, recurrence, pT (Amin et al., 2017), pN (Amin et al., 2017), pathologic stage (Amin et al., 2017), depth of invasion, histologic grade WHO (Barnes et al., 2005), invasion pattern, inflammatory response, degree of keratinization, nuclear pleomorphism, and perineural invasion (IBM SPSS Statistics 28.0 software; IBM Corp.; p ≤ 0.05; Student’s t-test or ANOVA).

#### Comparison between tumor and metastasis proteomes

To avoid deviations in protein identification and quantitation caused by searches in individual datasets, RAW data obtained from MS runs in malignant cells from the primary tumor and lymph nodes (n = 11 samples/each dataset) and for the proteome of non-malignant cells from both sites (n = 27 samples/each) were combined in a unique search for malignant cells and a single search for non-malignant cells using MaxQuant software as described in the ‘Discovery proteomics and data analysis” subsection. Log_2_ intensities were used to determine differentially abundant proteins between tumor and lymph nodes (p ≤ 0.05; Benjamini-Hochberg test). Differentially abundant proteins exhibiting higher expression in lymph nodes compared to that in primary sites or those exclusively detected in lymph nodes were termed “upregulated”, while proteins exhibiting lower expression in lymph nodes compared to that in primary sites or those exclusively detected in the tumor site were termed “downregulated”. GO biological processes were enriched using ShinyGO v. 0.61 (Enrichment FDR ≤ 0.05) (Ge et al., 2020). Hierarchical clustering of malignant and non-malignant populations was performed as reported in the ‘Clustering global proteome datasets’.

#### Reverse transcription quantitative PCR

Transcript levels of the genes *SRSF1, SRSF2, SRSF3, SRSF5, TRA2A,* and *CD209* were evaluated in a 19-patient cohort of lymph node tissues (9 pN+ and 10 pN0), a 24-patient cohort of buffy coats (10 pN+, 14 pN0), and a 22-patient cohort of saliva cells (10 pN+, 12 pN0), and they were then associated with nodal status. Total RNA was extracted using TRIzol reagent (Invitrogen, USA) and quantified using a Nanodrop ND-1000 spectrophotometer (NanoDrop Technology, USA). RNA was reverse transcribed using the High-Capacity cDNA Reverse Transcription Kit (Thermo Scientific, United States). PCR amplification was performed in an ABI Prism 7500 Sequence Detection System (Applied Biosystems, USA) using SYBR Mix (Applied Biosystems, USA). Primer set sequences were designed using the Primer-BLAST tool (http://www.ncbi.nlm.nih.gov/tools/primer-blast/), and IDT Oligoanalyzer^TM^ (http://www.idtdna.com/analyzer/applications/oligoanalyzer) was used to predict the occurrence of dimers and secondary structures **(Table S6-1)**. Serial dilutions (1:5) of a pool of cDNA samples were used to evaluate the amplification efficiency (E) according to the equation E = 10(−1/slope)−1. Primers possessing E values ranging from 95% to 105% were used for further analysis. PCR amplifications were performed in duplicate, and the specificity of the products was verified based on melting curve analysis. Negative and positive controls were included for each reaction. Positive controls consisted of a pool of samples representative of each biological material that was used as a ‘normalizing’ sample and allowed for the evaluation of intra-assay variations. *GAPDH* and *HPRT* were used as the reference genes. Relative quantification was calculated using the model proposed by Pfaffl (Pfaffl, 2001), and the Student’s t-test was applied to compare log_2_ transcript levels between pN+ and pN0 tissue samples (IBM SPSS Statistics 28.0 software, IBM Corp.; GraphPad Prism v8.2.1, GraphPad, https://www.graphpad.com; p ≤ 0.05).

#### Selection of targets for ML analysis

Using discovery proteomics, we determined sets of proteins in tissues and fluids from HNSCC patients that may be used as markers of lymph node metastasis. Thus, we defined a series of criteria to prioritize particular proteins for further verification using targeted methodologies, including (i) global proteome datasets with clustering patterns associated with nodal status, (ii) metastasis-associated datasets with the best segregation pattern using PCA, (iii) proteins detected exclusively in one group or with p ≤ 0.05 when using FDR correction to compare pN+ and pN0 groups based on the Benjamini-Hochberg test, (iv) high performance in discriminating patients according to nodal metastasis in ROC curve analysis (AUC > 0.7), (v) significant association with other clinical and pathological features (p ≤ 0.05; Student’s t-test, ANOVA or Fisher’s exact test), (vi) generation of high-performance protein or transcript pairs <Si, Cj> to separate pN+ and pN0 using ML (please see section “Definition of prognostic signatures using machine learning”), and (vii) association with the immune contexture highlighted by an elevated gene expression in lymphoid tissues compared to that of other tissue types according to The Human Protein Atlas database (list of 1,419 elevated genes in the lymphoid tissue transcriptome available from http://www.proteinatlas.org) **(data not shown)**. Six microenvironment proteins satisfying most (if not all) of the criteria were selected for verification in buffy coat and saliva samples using SRM-MS and RT-qPCR for SRSF1, SRSF2, SRSF3, SRSF5, TRA2A (fulfilling criteria [i] to [vi]), and CD209 (fulfilling criteria [i] to [iii] and [vi] to [vii]). Moreover, due to the relevance of the immune system highlighted in several analyses throughout this work, 19 immune markers were selected for assessment in fluids using SRM-MS to quantify the proteins associated with immune populations, including leukocytes (CD45), myeloid cells (CD11b), T lymphocytes (CD3), CD4+ T lymphocytes (CD4), CD8+ T lymphocytes (CD8), Tregs (FOXP3 and CD25), B lymphocytes (CD19), NK cells (CD56 and CD16), monocytes (CD14, and CD64), non-classical monocytes (CD16), macrophages (CD64 and CD163), neutrophils (CD16, CD66b, and CD15), dendritic cells (CD11c and HLADR) and activated cells (CD80 and CD86).

#### Selection of proteotypic peptides and transitions for targeted proteomics

Proteotypic peptides were selected for each of the 26 microenvironments or immune targets based on the number of residues, their hydrophobicity, the presence in our DDA datasets from lymph nodes (non-malignant cells), buffy coat, or saliva, and SRM-Atlas evidence when the peptide was not identified in our data (Gallien et al., 2011; Lange et al., 2008). The selection of the best transitions was performed using spectral libraries built from our own DDA data, and transitions retrieved from the SRMAtlas were included when they were not detected with high reliability from DDA. The most intense transitions were selected for the final SRM-MS method. To monitor the microenvironment proteins associated with lymph node metastasis (SRSF1, SRSF3, SRSF5, TRA2A, and CD209), nine proteotypic peptides were selected and purchased as crude heavy-isotope-labeled peptide standards (JPT Peptide Technologies, USA) **(Table S6-2)**. SRSF2 did not possess proteotypic peptides fulfilling the criteria mentioned above and was excluded from the SRM-MS analysis. Stable isotope-labeled peptides were synthesized with ^13^C_6_,^15^N_2_-lysine or ^13^C_6_,^15^N_4_-arginine (+8 or +10 Da, respectively) that were localized preferentially at the C-terminal of the peptide. Three to four transitions were monitored in light and heavy channels for each peptide for a total of 68 transitions **(Table S6-3)**. The concentration of heavy peptides was optimized based on serial dilutions in buffy coat (1:2 and 1:4 dilutions) and saliva (1:2, 1:10, 1:30, and 1:60 dilutions) samples to determine the best concentration with a maximum 1:10 light/heavy ratio, and this resulted in a range of 9 to 200 fmol/µg of matrix. Considering the 19 immune markers (CD45, CD3, CD4, CD8, FOXP3, CD25, CD19, CD56, CD16, CD11b, CD14, CD64, CD163, CD66b, CD15, CD11c, HLADR, CD80, and CD86), heavy peptides were not included in the analysis, and the retention times were predicted by building a calculator in Skyline 19.1 software (MacLean et al., 2010) based on the spiked iRT mixture (Pierce™ Peptide Retention Time Calibration Mixture, Thermo Scientific, USA) while considering spectral libraries built from lymph node FFPE tissues, buffy coat, and saliva DDA data. When the calculator could not be applied due to the lack of peptide detection in our DDA libraries, transitions available from the SRMAtlas were monitored. Two to four peptides were included for each of the 19 immune proteins, and a total of 54 label-free peptides were monitored **(Table S6-2)**. Of the 19 targets, nine were detected in the DDA dataset **(Table S6-2).** None of the peptides from 10 out of 19 markers matched the retention times predicted from the calculator or the transitions described in the SRMAtlas in one buffy coat and/or one saliva sample from HNSCC patients, and they were excluded from further analysis. The remaining nine immune markers that included CD3 (T lymphocytes), CD4 (CD4+ lymphocytes), CD8 (CD8+ lymphocytes), CD11b (myeloid cells), CD14 (monocytes), CD16 (neutrophils, NK, non-classic monocytes), CD19 (B lymphocytes), CD45 (leukocytes), and CD66b (neutrophils) were retained for subsequent analysis, and four to six transitions were monitored in light channels per peptide for a total of 80 transitions **(Table S6-3)**.

#### Targeted proteomics and data analysis

The relative abundance of six prioritized microenvironment proteins from lymph nodes (SRSF1, SRSF2, SRSF3, SRSF5, TRA2A, and CD209) and nine immune markers (CD3, CD4, CD8, CD11b, CD14, CD16, CD19, CD45, and CD66b) was quantitatively evaluated in 19 buffy coat (7 pN+, 12 pN0) and 25 saliva (16 pN+, 9 pN0) samples from HNSCC patients using SRM-MS targeted proteomics. Sample preparation was performed as described in the “Sample preparation for discovery proteomics” subitem. Samples were analyzed on a Xevo TQ-XS triple quadrupole mass spectrometer (Waters, USA) as described previously (Carnielli et al., 2018; Neves et al., 2020). Buffy coat or saliva digests (1.6 µg) were resolved over a 60-min gradient using an Acquity UPLC-Class M equipped with a trap column (Acquity UPLC BEH C18 130A, 5 μm, 300 μm × 50 mm, Waters, USA) and a BEH Shield C18 IonKey column (10-cm × 150-μm ID packed with 1.7-μm C18 particles, Waters, USA) at 1.2 a flow rate and a temperature of 40 °C. The MeCN gradient started at 2% B (MeCN, 0.1% formic acid), and this was followed by a linear ramp to 40% B over 45 min, then by a step increase to 85% B for 47 min, and finally conditioning at 2% B for 60 min. Mass spectrometry analysis of eluting peptides was performed via SRM-MS mode with quadrupoles Q1 and Q3 operating as unit mass resolution (0.7 Th full width at half maximum). The optimal collision energy was determined for each peptide using Skyline 19.1 (MacLean et al., 2010). Scheduled SRM-MS acquisition was adjusted to a 3-min elution window with dwell times automatically set in MassLynx v4.2 to achieve at least ten points per peak over a 30-s elution profile. A blank sample (water) was injected between two consecutive sample runs with the same gradient to minimize carryover. Peaks were inspected using Skyline 19.1 (MacLean et al., 2010) and quantified by calculating the ratio between light and heavy intensities for each peptide (intensity = sum of transitions). For immune markers, light intensities were considered for quantification.

#### Quality control in targeted proteomics

Several quality controls were used to evaluate the reliability of the SRM-MS analysis. The percentage of carryover was assessed by comparing the absolute intensity of 13 heavy peptides **(Figure S5A)** in a blank (water) injection with the absolute intensity of the same peptides in a preceding matrix (buffy coat) run (Abbatiello et al., 2013). A system suitability protocol was implemented to assess equipment performance by monitoring 18 peptides from a mixture of 200 ng of digested bovine serum albumin (BSA) and 5 fmol of iRT (Pierce™ Peptide Retention Time Calibration Mixture, Thermo Scientific, USA) prior to each batch of four samples. To monitor sample quality, an iRT mixture (32 fmol injected) was spiked into each sample, and four peptides with their respective three or four transitions each were monitored as a control for retention time and intensity shifts in liquid chromatography **(Figure S5B-E; Table S6-2)**. The coefficient of variation (CV) was calculated, and samples exhibiting a CV ≥ 20% were manually evaluated. Moreover, both light and heavy peptides from SRSF1, SRSF3, SRSF5, TRA2A and CD209 were assessed in Skyline 19.1 (MacLean et al., 2010) by observing the alignment of elution times, co-elution of all transitions, relative intensity correlation with the spectral library (dotp, close to 1), and proximity to the predicted retention time. The relative intensity correlation between light and heavy transitions (rdotp) was also evaluated, resulting in (i) the inclusion of peptides from samples exhibiting rdotp ≥ 0.9, (ii) exclusion of peptides possessing rdotp ≤ 0.8, and (iii) manual evaluation of peptides exhibiting rdotp varying from 0.8 to 0.9. For immune markers, peaks for light peptides were evaluated for the predicted elution time and co-elution of all transitions.

#### Cross validation of protein and gene expression

RNASeq samples of 500 HNSCC primary tumor samples were retrieved from TCGA (https://portal.gdc.cancer.gov) using the TCGA-HNSC identifier. Gene expression was compared to the protein levels in malignant cells from the primary tumor. Spearman correlations using the median FPKM from HNSCC were performed using the mean LFQs of proteomes.

#### scRNASeq processing and differential expression

The HNSCC scRNASeq dataset from 18 HNSCC patients was retrieved from the Gene Expression Omnibus using the query number GSE103322 (Puram et al., 2017). We followed the standard workflow from Seurat package v.4.0.0 (Hao et al., 2020) with parameters min.cells=0 and min.features=200 for matrix import and nfeatures = 2000 for FindVariableFeatures(), and 30 dimensions were used after reduction with principal component analysis (PCA) followed by uniform manifold approximation and projection (UMAP) map construction. Clusters were identified using a resolution of 1.2 and the cell types were annotated according to the literature (Puram et al., 2017). Additional annotation was made to subdivide fibroblasts into CAFs (expressing *FAP* and *THY1*), myofibroblasts (expressing *ACTA2*), and skeletal muscle cells (expressing *DES*), and to identify T cell subtypes. The original annotation of plasma/B cells was curated and updated based on B cell (*MS4A1* and *BANK1*) and plasma cell markers (*MZB1* and *IGLL5*). For each population within the HNSCC dataset, markers that define clusters via differential expression (cluster markers) from previous analysis were tested for differential expression among subpopulations using Wilcox, MAST, and Bimod tests within the FindMarkers() function and “BH” as p adjustment method. The PBMC 3k dataset was retrieved at https://support.10xgenomics.com/single-cell-gene-expression/datasets/1.1.0/pbmc3k and processed as previously reported (Hao et al., 2020). Cluster markers in PBMC populations were identified using the FindAllMarkers function with min.pct = 0.1 and logfc. threshold = 0.25. Differentially expressed proteins from the buffy coat, tumor microenvironment, and lymph node microenvironment were selected and used to query filtered marker genes (adjusted p 0.05). The difference in LFQ proteome abundance between the pN+ and pN0 groups was calculated prior to plotting.

#### Cell type deconvolution and enrichment

Cell type annotations provided in the processed matrix from the literature (Puram et al., 2017) were used to generate a signature matrix specific for HNSCC to infer cell types in bulk proteome and transcriptome data. The signature matrix was created using the following parameters: Min. expression = 0, replicates = 4, and sampling = 1. Proteomic data (LFQ intensities) from primary HNSCC samples were used to infer cell type abundance from the scRNASeq signature matrix using CIBERSORTx with default parameters and 100 permutations (Newman et al., 2019). The cell signature derived from healthy PBMC subpopulations was used as a reference to deconvolute whole blood proteome information using LFQ intensities. For deconvolution, data from both pN0 and pN+ samples were used, and a deconvolution (p ≤ 0.01) was set to determine significance (Steen et al., 2020).

xCell, which performs Gene-set enrichment analysis in previously defined immune and stromal cell types, was used to evaluate differences between pN+ and pN0 samples (Aran et al., 2017). TCGA data available on the xCell website (https://xcell.ucsf.edu/) were downloaded and filtered using clinical information from cBioportal as Sample.Type == “Primary” and excluding samples without lymph node stage annotation. HNSCC pN+ and pN0 samples (n = 428 patients) were classified according to information in “Neoplasm.Disease.Lymph.Node.Stage.American.Joint.Committee.on.Cancer.Code.” A subset of oral cavity samples (OSCC, n = 248 patients) was also analyzed using clinical data from cBioportal. Spearman correlations were performed between the signature scores and pN+/pN0 outcome filtering for p ≤ 0.05.

#### RNASeq data analysis

Bioinformatics analysis of third-party RNAseq data was performed in R environment (v4.0.0) using Hmisc v4.4-0 for correlations, dplyr v1.0.0, tibble v3.0.1, data.table v1.13.0, and readxl v1.3.1 for data manipulation, SummarizedExperiment v1.18.2, GEOquery v2.56.0 (Davis and Meltzer, 2007), and TCGAbiolinks v2.16.0 (Colaprico et al., 2016) for data retrieval, ggpubr v0.3.0, ggplot2 v3.3 for plotting, and Seurat v4.0.0 (Hao et al., 2020) for scRNASeq analysis and manipulation.

#### Definition of prognostic signatures using machine learning

The predictive power of peptides, proteins, and/or transcripts from the microenvironment (SRSF1, SRSF2, SRSF3, SRSF5, TRA2A, CD209) and immune targets (CD3, CD4, CD8, CD11b, CD14, CD16, CD19, CD45, CD66b) that were selected as described above and their combination to distinguish HNSCC patients based on locoregional metastasis status was determined using an ML approach (pN+ vs. pN0). The steps used to obtain and validate the signatures are presented in **Figure 7A**. First, (1) quantitation data acquired for the microenvironment and immune targets in buffy coat and saliva using SRM-MS or RT-qPCR were selected (individual datasets: n = 19 and 24 patients for SRM-MS and RT-qPCR in buffy coat, respectively; n = 25 and 22 patients for SRM-MS and RT-qPCR in saliva, respectively; combined datasets: n = 15 patients for buffy coat, n = 14 patients for saliva). We also performed ML analysis of protein (DDA) and transcript (RT-qPCR) data from lymph node microenvironment cells with the purpose of selecting targets (please see section “Selection of targets for verification”). In total, we used 14 datasets representing the quantification of peptides, proteins, transcripts, and their combinations. The missing values were replaced with 0. For each dataset, we ran a sequence of steps to capture a set of candidate signatures that performed well in the classification task (pN+ vs. pN0). Then, (2) variables were combined to create all possible signatures (Si) of size 1 to N, where N is the maximum number of peptides/proteins/transcripts in each signature and was arbitrarily set to 5. Next, (3) each signature was used to create different classification models (Cj) that included ridge, linear SVM, lasso, linear discriminant, perceptron, decision tree, naive Bayes, GBM, random forest, and RBF SVM. Each pair signature-classifier <Si, Cj> was evaluated within a repeated stratified k-fold cross-validation. The total number of tested models per pair <Si, Cj> was defined as R * K, with R = 10 repetitions and K = smallest class size. Finally, (4) a list of potential signatures <Si, Cj> to discriminate pN+ and pN0 patients was selected for each dataset considering the evaluation step (3). The pair <Si, Cj> with the highest average ROC AUC was named ‘top-1’, and the respective curves were represented. Furthermore, we identified pairs <Si, Cj> with potential equivalent performance, comprising the pairs for which the distribution of AUC values observed in the repeated cross-validation was not statistically different from the ones observed with the top-1 pair (p ≥ 0.05; Student’s t-test). Specifically, if we could not reject the hypothesis of a pair P possessing an equal average ROC AUC to the Top-1 pair, we included pair P in the final list of selected pairs. Top-1 and all selected pairs were characterized as high-performance signatures. In addition to AUC, sensitivity, specificity, and precision were recorded for every high-performance pair <Si, Cj>. Heatmaps were generated using GraphPad Prism v8.2.1 (GraphPad software; https://www.graphpad.com).

#### Flow cytometry analysis

The expression of SRSF3 and TRA2A proteins was evaluated in a 20-HNSCC patient cohort (10 N+, 10 N0) using flow cytometry. Cell suspensions were stained for surface markers using anti-CD45-BUV805, anti-CD3-BV421, anti-CD4-BV605, anti-CD8-BV650, anti-CD25-BUV563, anti-CD56-APC, anti-CD19-BV750, anti-CD209-PE-Cy7, anti-CD14-APC-Cy7, anti-CD15-PE and anti-CD11b-BV650 monoclonal antibodies by incubation for 30 min with antibody solutions, followed by washes. TRA2A antibody was labeled by Zenon™ Alexa Fluor™ 750 Rabbit IgG Labeling Kit (Thermo Scientific, United States), as recommended by the manufacturer. The cells were permeabilized using the BD Pharmingen™ Transcription Factor Buffer Set (BD Biosciences, United States) for 40 min at 4°C and further stained with anti-SRSF3-FITC and anti-TRA2A-Alexa Fluor 750 for 40 min at 4°C followed by washes. Two PBMC samples from HNSCC patients were included as controls for surface markers. The preparations were analyzed using a FACSymphony^TM^ equipment (BD Biosciences, United States) and the FlowJo v10.8 software (BD Biosciences, United States). At least 30,000 gated events were acquired assuring the reliability of positive populations. Comparisons between N+ and N0 groups were performed using MFI and proportion of cells in the GraphPad Prism v8.2.1 software (GraphPad; https://www.graphpad.com). Data were tested for normality and homogeneity of variance using Shapiro-Wilk test to further guide the selection of the appropriate statistical test used to verify differences between N+ and N0 samples (Student’s t-test, Welch’s t-test or Mann-Whitney tests; p ≤ 0.05).

### QUANTIFICATION AND STATISTICAL ANALYSIS

Statistical analyses were performed using R environment v4.0.0, IBM SPSS Statistics 28.0 (IBM Corp.) or GraphPad Prism v8.2.1 (GraphPad; https://www.graphpad.com) and are indicated in the figure legends, results, and methods sections. All proteomics quantitative datasets were log_2_ transformed to reduce skewness, and a parametric Student’s t-test was used for group comparison. Multiple comparisons were adjusted using the Benjamini-Hochberg correction when feasible. The association between protein abundance and clinicopathological characteristics was verified using Student’s t-test, ANOVA, or Fisher’s exact test. Flow cytometry data were analyzed considering the appropriate parametric or non-parametric test (Student’s t-test, Welch’s t-test or Mann-Whitney test). Statistical significance was established at p ≤ 0.05. To prevent bias during DDA and SRM-MS measurements, tissue and fluid samples were randomized in the R environment **(Tables S2-1 and S5-1)**. Two samples were excluded from MS analysis due to deviations in quality control measurements (malignant cells from primary tumors 4417 and 2875) **(Figure S1)**.

## SUPPLEMENTARY INFORMATION TITLES AND LEGENDS

### Supplementary Figures

**Figure S1. iRT and trypsin quality control for tissue and fluid samples evaluated by mass spectrometry (LC-MS/MS, DDA), Related to Figure 1**.

(A) Intensities and retention times for 4 iRT peptides (50 fmol) used to monitor deviations in mass spectrometry runs for 94 individual FFPE samples and for each group of comparisons. Black circles indicate discrepant intensities for the four iRT peptides in case 66 (malignant cells from primary tumor 4417), and the arrows show absent intensity and retention time for iRT_Pep4 in case 71 (malignant cells from primary tumor 2875). The two samples were excluded from further analysis due to the inconsistent profile. No deviations were observed in the chromatographic pattern among groups. FFPE: Formalin-Fixed Paraffin-Embedded tissue sample, PT_M: primary tumor – malignant cells, PT_NM: primary tumor – non-malignant cells, LN_M: lymph node – malignant cells, LN_NM: lymph node – non-malignant cells.

(B) Retention times for 3 trypsin autolysis peaks (*m*/*z* 421.7584, +2; *m*/*z* 523.2855, +2; *m*/*z* 737.7062, +3) used to monitor deviations in sample preparation and LC-mass spectrometry runs for 94 tissues, 24 buffy coats, and 24 saliva samples. Black arrows indicate sample 71 that was missing peaks for the three trypsin peptides evaluated. The sample was excluded from proteomics analysis (sample 2875 – malignant cells from primary tumor). FFPE: Formalin-Fixed Paraffin-Embedded tissue sample.

**Figure S2. Characterization of tissues and fluids according to the proteomic profile in a 59-HNSCC cohort, Related to Figure 1**.

(A) Upset plot presenting shared and exclusive proteins for the 6 HNSCC sites that were evaluated. Proteomics data acquired for each site were run independently in MaxQuant software.

(B) Heat maps revealing the proteins clusters (PC1, PC2 and/or PC3) that were identified for cell populations from tissues and fluids. PC groups were generated using the following methods and distances: Ward Sqeuclidean (25 primary tumor – malignant: 2,451 proteins), Weighted Camberra (27 primary tumor – non-malignant: 1,984 proteins and 24 buffy coats: 2,188 proteins), Ward Sqeuclidean (13 lymph node – malignant: 2,374 proteins and 27 lymph node – non-malignant: 2,137 proteins), and Complete Sqeuclidean (24 saliva samples: 1,154 proteins).

**Figure S3. Association of the proteome landscape with immunity and clinical and pathological data, Related to Figures 3, 5 and 6**.

(A) Spearman correlation coefficients (R) generated by comparing the proteome composition from HNSCC malignant cells from our dataset to the RNASeq data of tumors from TCGA.

(B-C) Annotation of populations from HNSCC primary tumors and the lymph node microenvironment of our dataset in the HNSCC scRNASeq dataset (Puram et al., 2017). The standard workflow from Seurat was used to cluster cells. Annotations from the literature (Puram et al., 2017) were used as guide to further divide fibroblasts into CAFs and myofibroblasts. (C) shows expression of cluster-defining genes used for annotations in (B).

(D) Spearman correlation coefficients (R) generated for the comparison of nodal status (last column) and immune populations. RNASeq data from OSCC patients from TCGA with information of nodal metastasis was used to identify cell signatures using the xCell algorithm. Correlations with p ≤ 0.05 are presented.

(E) Average fold change and differential expression of 6 targets selected for verification in non-immune populations from the tumor microenvironments (pN+ vs. pN0). The dashed lines represent a p threshold of 0.05 and significant results are labeled with empty circles.

(F) Clinical-pathological features from HNSCC patients that were significantly associated with patient clusters C1 and C2 for tissues (p ≤ 0.05; Fisher’s exact test). *p ≤ 0.05.

(G) Clinical-pathological features from HNSCC patients that were significantly associated with patient clusters C1 and C2 for buffy coat and saliva samples. (p ≤ 0.05; Fisher’s exact test or log-rank test). *p ≤ 0.05.

(H) Evaluation of HPV DNA and RNA in primary tumor tissues (n = 27 samples) from HNSCC patients. NA: not available.

**Figure S4. Characterization of nodal metastasis biomarkers detected in the lymph node microenvironment from 27 HNSCC patients, Related to Figure 5**.

(A) Clinical-pathological features significantly associated with the abundance of SRSF1, SRSF2, SRSF3, SRSF5, and TRA2A proteins (p ≤ 0.05; Student’s t-test, ANOVA, and Fisher’s exact test). High and low levels of proteins were defined using the mean intensity as the cut-off. WD: well differentiated tumor, MD: moderately differentiated tumor, PD: poorly differentiated tumor, AUC: area under the curve. *p ≤ 0.05, **p ≤ 0.01, ***p ≤ 0.001.

(B) Protein interaction networks for the proteins associated with lymph node metastasis.

(C) LFQ log_2_ intensity for SRSF1, SRSF2, SRSF3, SRSF5, TRA2A, and CD209 in lymph node tissues from HNSCC patients as determined by DDA (pN+ vs pN0; q ≤ 0.05; Benjamini-Hochberg test). CD209 was detected only in pN0 samples. *q ≤ 0.05, **q ≤ 0.01.

(D) Relative expression of *SRSF1, SRSF2, SRSF3, SRSF5, TRA2A,* and *CD209* in lymph node tissues from HNSCC patients as determined by RT-qPCR (pN+ vs pN0; p ≤ 0.05; Student’s t-test). *p ≤ 0.05, **p ≤ 0.01, ***p ≤ 0.001.

**Figure S5. Quality control of SRM-MS experiments, Related to Figure 7**.

(A) Frequency of carryover for heavy peptides from the microenvironment biomarkers SRSF1, SRSF3, SRSF5, TRA2A, and CD209 (14.4 to 320 fmol) and for 4 iRT peptides (32 fmol) (peptide intensity in water/peptide intensity in buffy coat matrix).

(B-C) Peak areas for iRT mixture (32 fmol) obtained from mass spectrometry runs in buffy coat (B) and saliva (C) samples.

(D-E) Retention time for iRT mixture (32 fmol) obtained from mass spectrometry runs in buffy coat (D) and saliva (E) samples.

**Figure S6. Transitions monitored by SRM-MS for all targeted peptides, Related to Figure 7**.

(A-B) Representative peaks indicating transitions monitored for the immune (CD3, CD4, CD8, CD11b, CD14, CD16, CD19, CD45, and CD66b) (A) and microenvironment (SRSF1, SRSF3, SRSF5, TRA2A, and CD209) (B) markers selected for SRM-MS analysis **(Table S6-2 and 3)**. Predicted retention times are presented for the immune peptides (when available).

**Figure S7. Characterization of the highest-performance signatures according to machine learning analysis, Related to Figure 7**.

(A-B) ROC curves indicating the top-1 pairs <Si, Cj> for peptides, proteins, or transcripts from individual microenvironments and immune datasets from buffy coat (A) and saliva (B). The top-1 pair possessing the higher AUC across datasets is presented in Figure 7D. AUC: area under the curve.

(C-D) Count of microenvironment and immune targets in signatures from buffy coat samples (C) and saliva (D) cells for the combined analysis of peptides, proteins, and transcripts datasets. Only signatures exhibiting high performance for discriminating pN+ and pN0 HNSCC patients were considered (top-1 pair <Si, Cj> and signatures with p ≥ 0.05; Student’s t-test).

(E) Flow cytometry gating strategy for immune populations from buffy coat samples. Total leukocytes were first gated on a side scatter (SSC-A)/CD45 plot to define lymphocytes and myeloid cells (upper left). The lymphocytes were then gated on the CD19+ (B cells), CD56+CD3- (NK cells) and CD56-CD3+ (T lymphocytes) populations (left). These were further gated on the CD4+, CD8+ and CD4+CD25+ (activated T lymphocytes) subsets (left). The myeloid cells were separated by CD11b expression and further phenotyped according to CD209 (dendritic cells), CD15 (neutrophils) and CD14 (monocytes) expression (right).

(F) Histogram of the flow cytometry analysis showing the expression of SRSF3 and TRA2A in buffy coat samples from all N+ and N0 HNSCC samples (10 N+; 10 N0).

## Supplementary Tables

**Table S1.** Samples and main clinical features of HNSCC patients included in this study, Related to Figure 1.

**Table S2.** All proteins identified across multiple sites after excluding reverse sequences *(Data based on MaxQuant and Perseus analyses),* Related to Figures 1 and 4.

**Table S3.** Differentially abundant proteins between pN+ and pN0 samples across multiple sites *(Data based on MaxQuant and Perseus analyses),* Related to Figures 2 and 4.

**Table S4.** Metastasis signatures using machine learning in protein and transcript datasets from multiple sites, Related to Figures 5 and 7.

**Table S5.** SRM-MS and RT-qPCR data for all peptides or transcripts quantified in buffy coat and saliva samples, Related to Figure 7.

**Table S6.** Oligonucleotides and peptides selected for RT-qPCR and SRM-MS experiments, respectively, Related to STAR Methods.

**Table S7.** Flow cytometry analysis of SRSF3 and TRA2A proteins in buffy coat samples from HNSCC patients, Related to Figure 7.

## Highlights

1. A deep mass spectrometry-based proteomics profiles multiple HNSCC sites
2. Immune modulation is overrepresented in HNSCC multisites
3. Malignant and non-malignant proteomes show different behavior towards tumor spread
4. Multiparametric machine learning reveals high-performance metastasis signatures

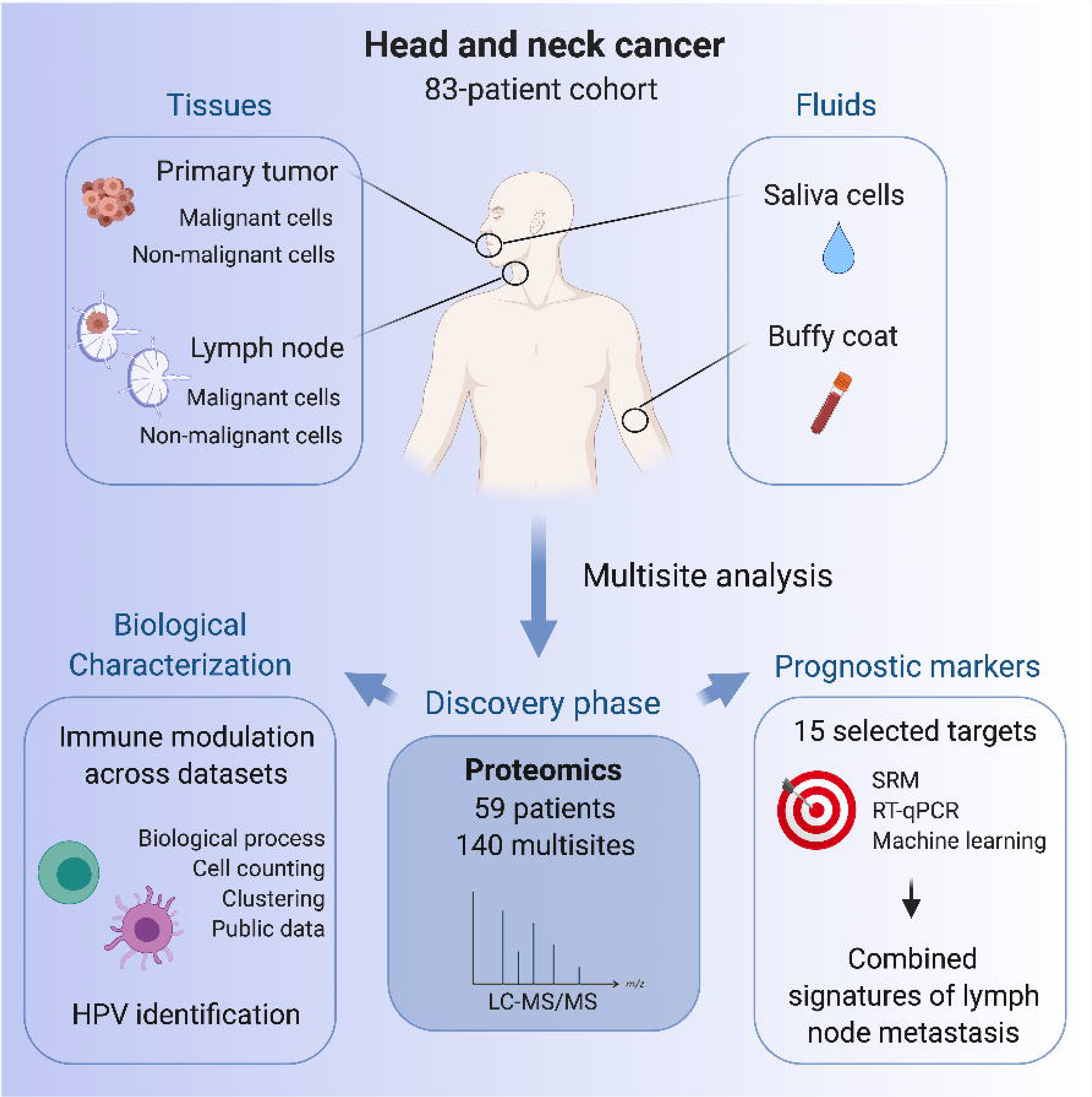

## Notes

### Competing Interest Statement

The authors have declared no competing interest.

