## Supplementary figures and images for "Wiring multiple microenvironment proteomes uncovers the biology in head and neck cancer"

### Figure S1

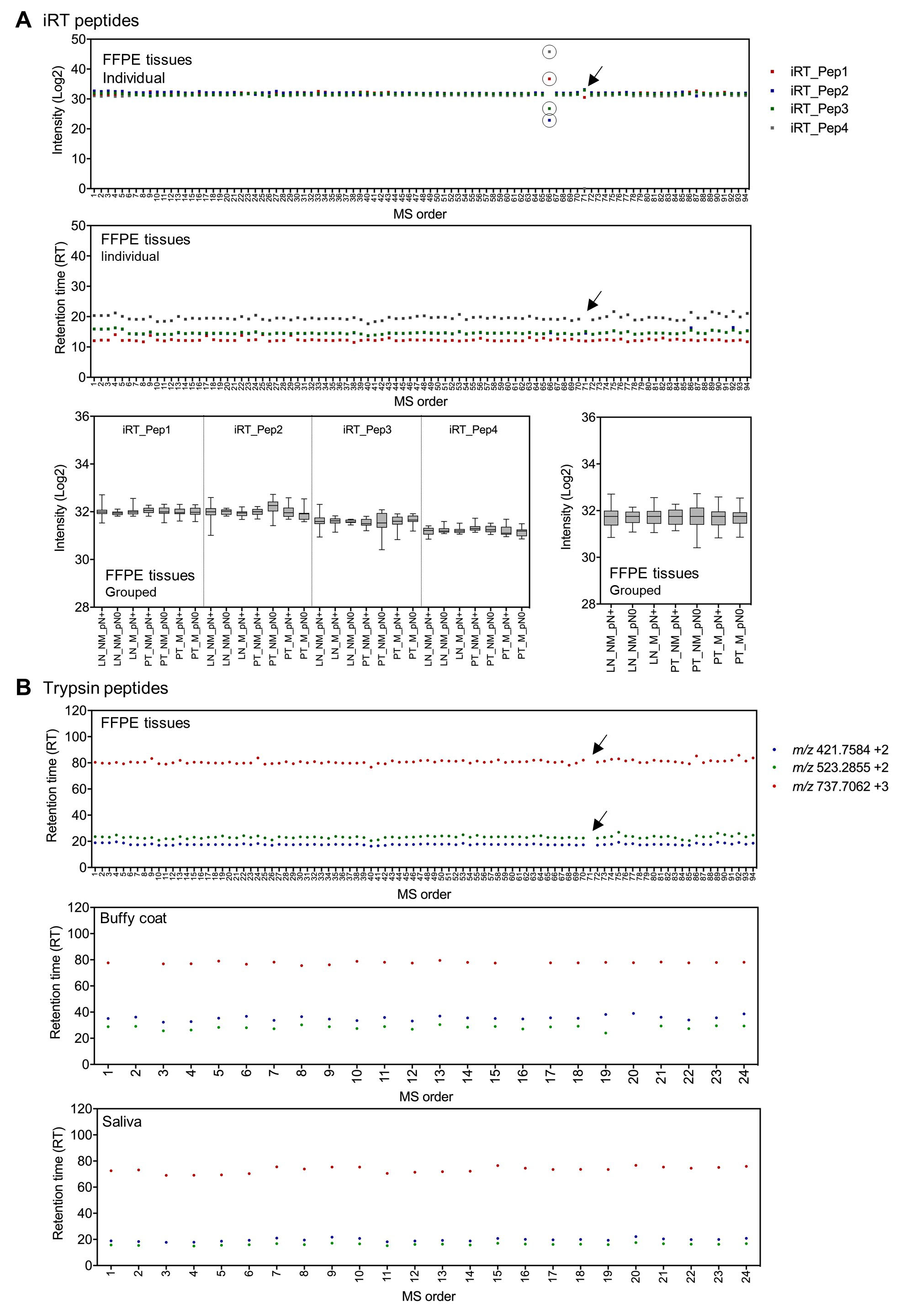

### Figure S2

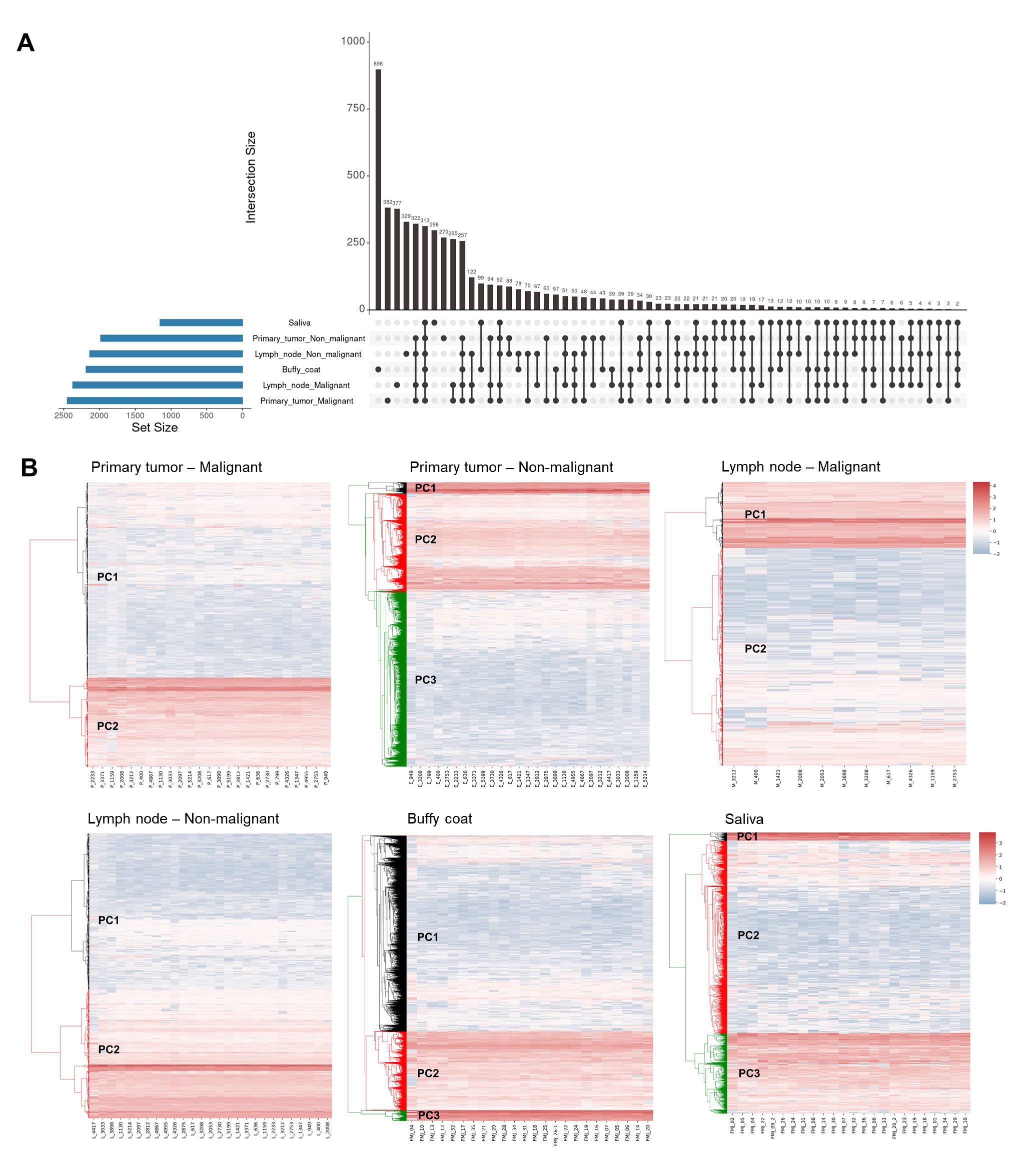

### Figure S3

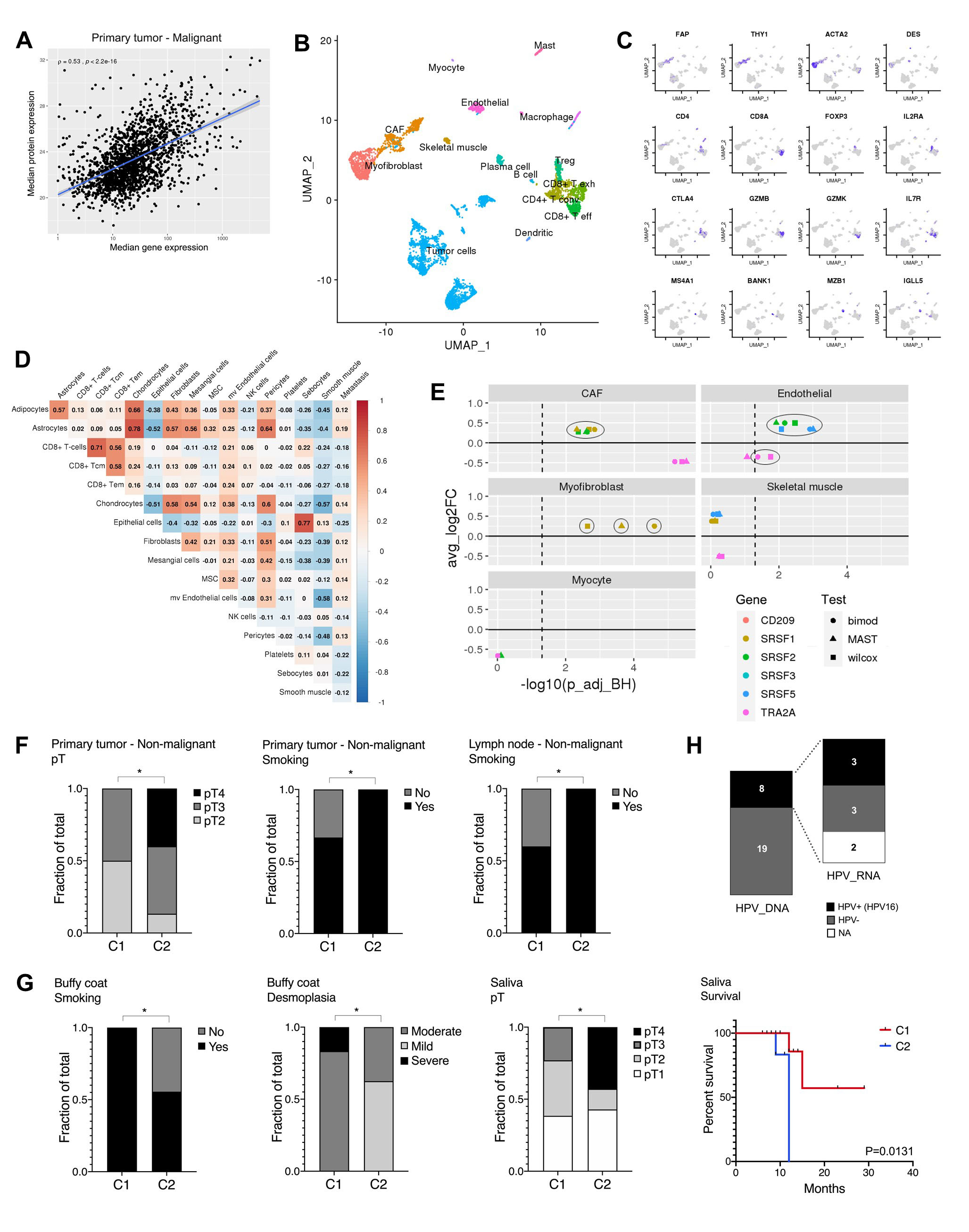

### Figure S4

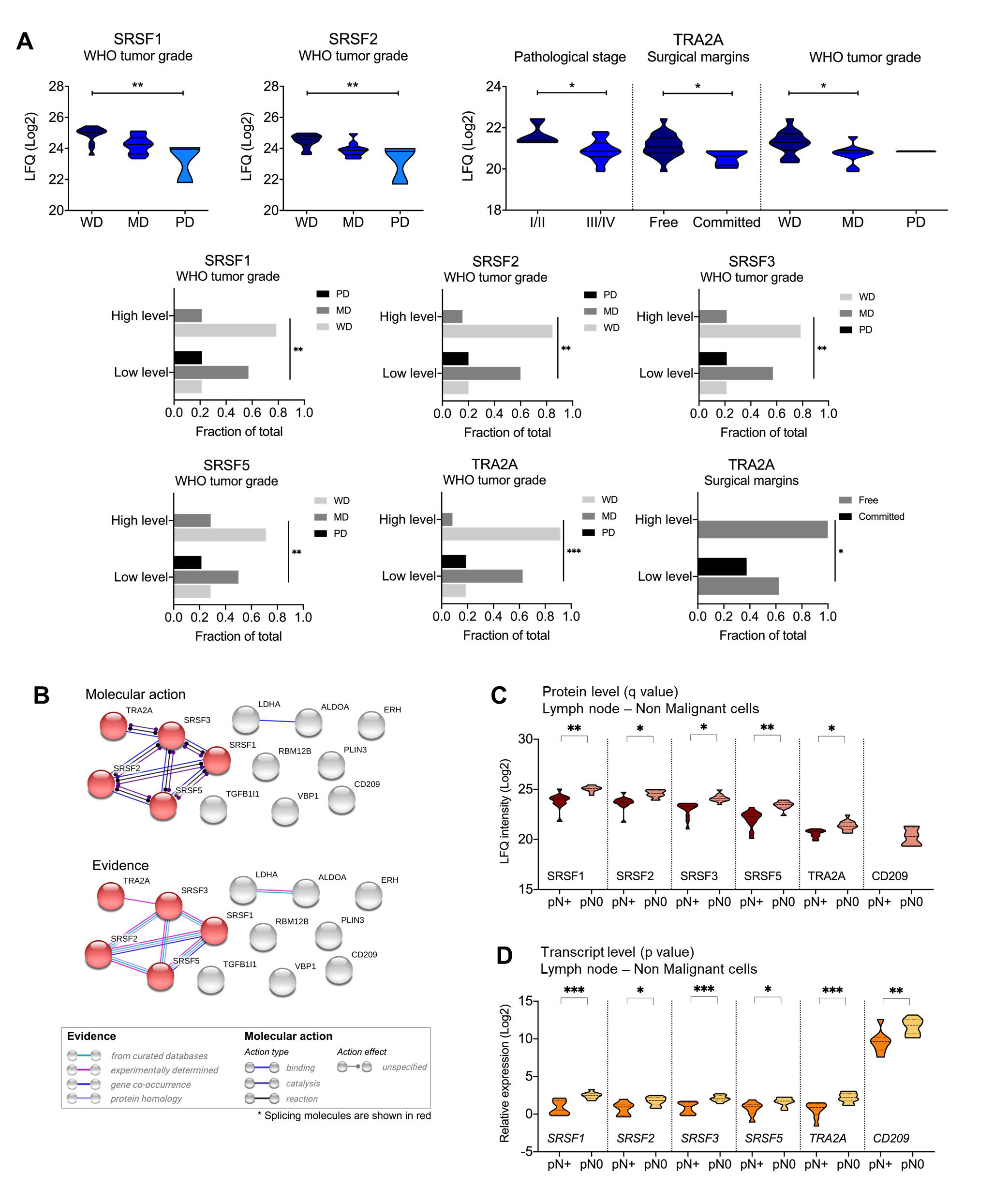

### Figure S5

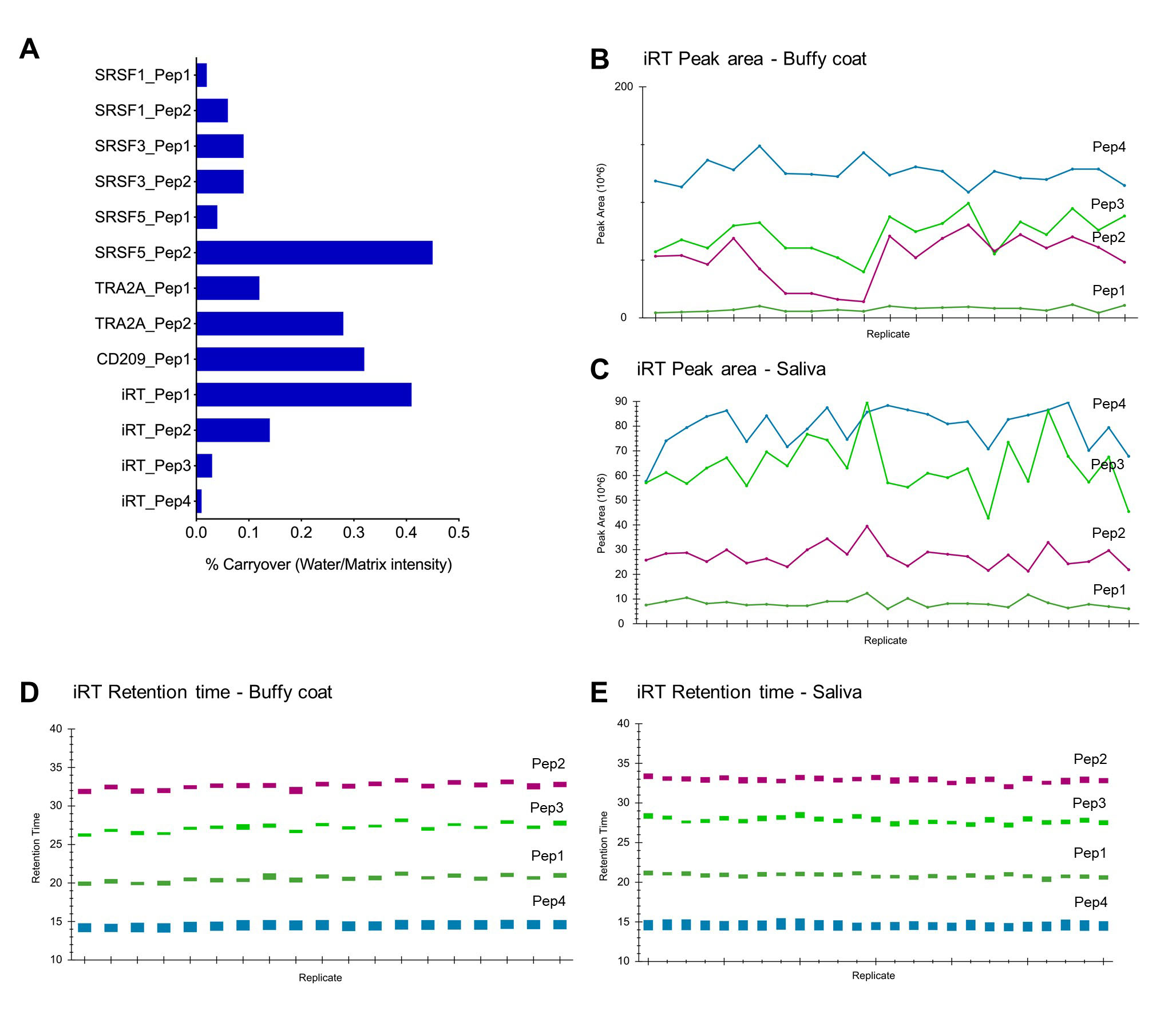

### Figure S6

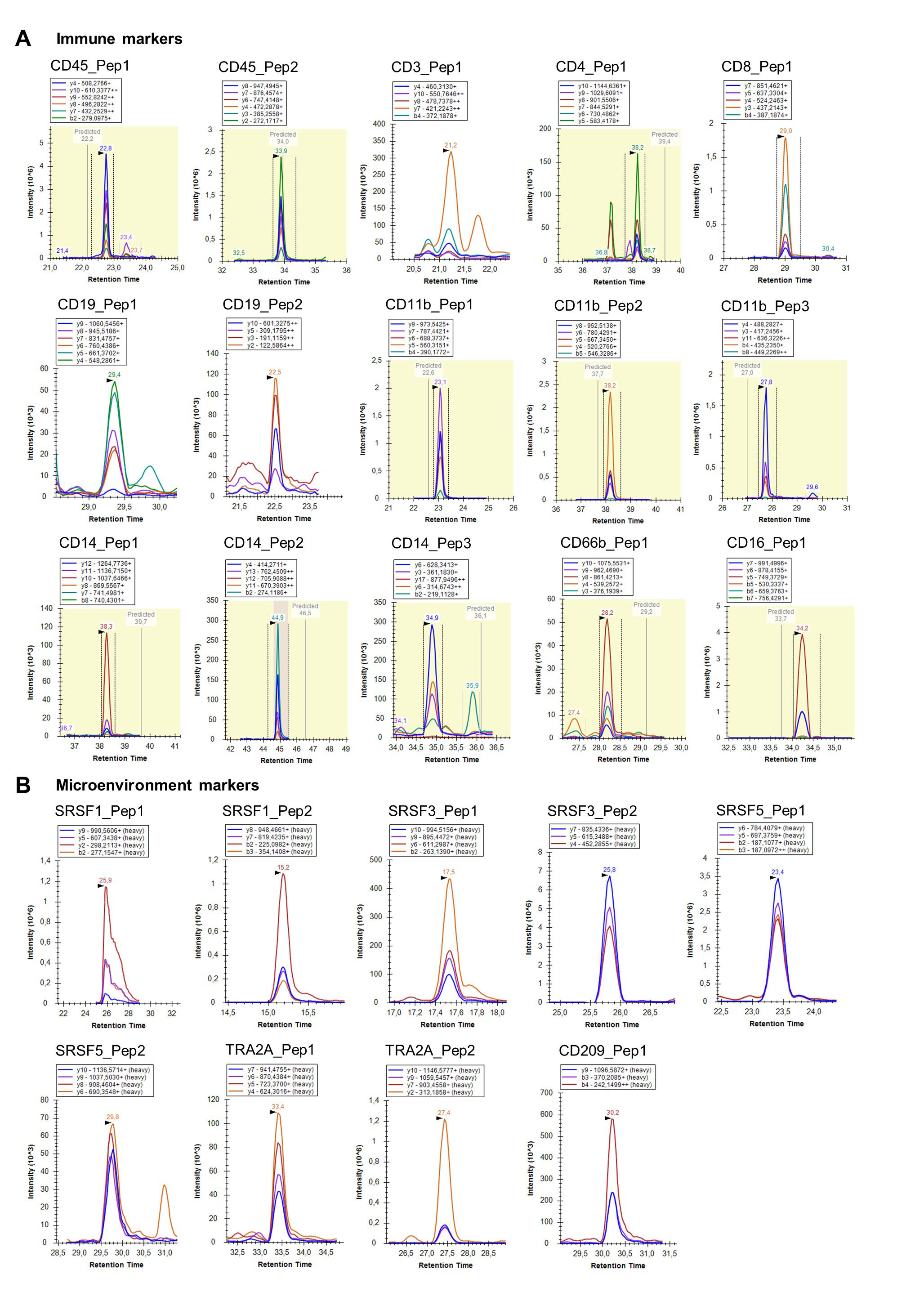

### Figure S7

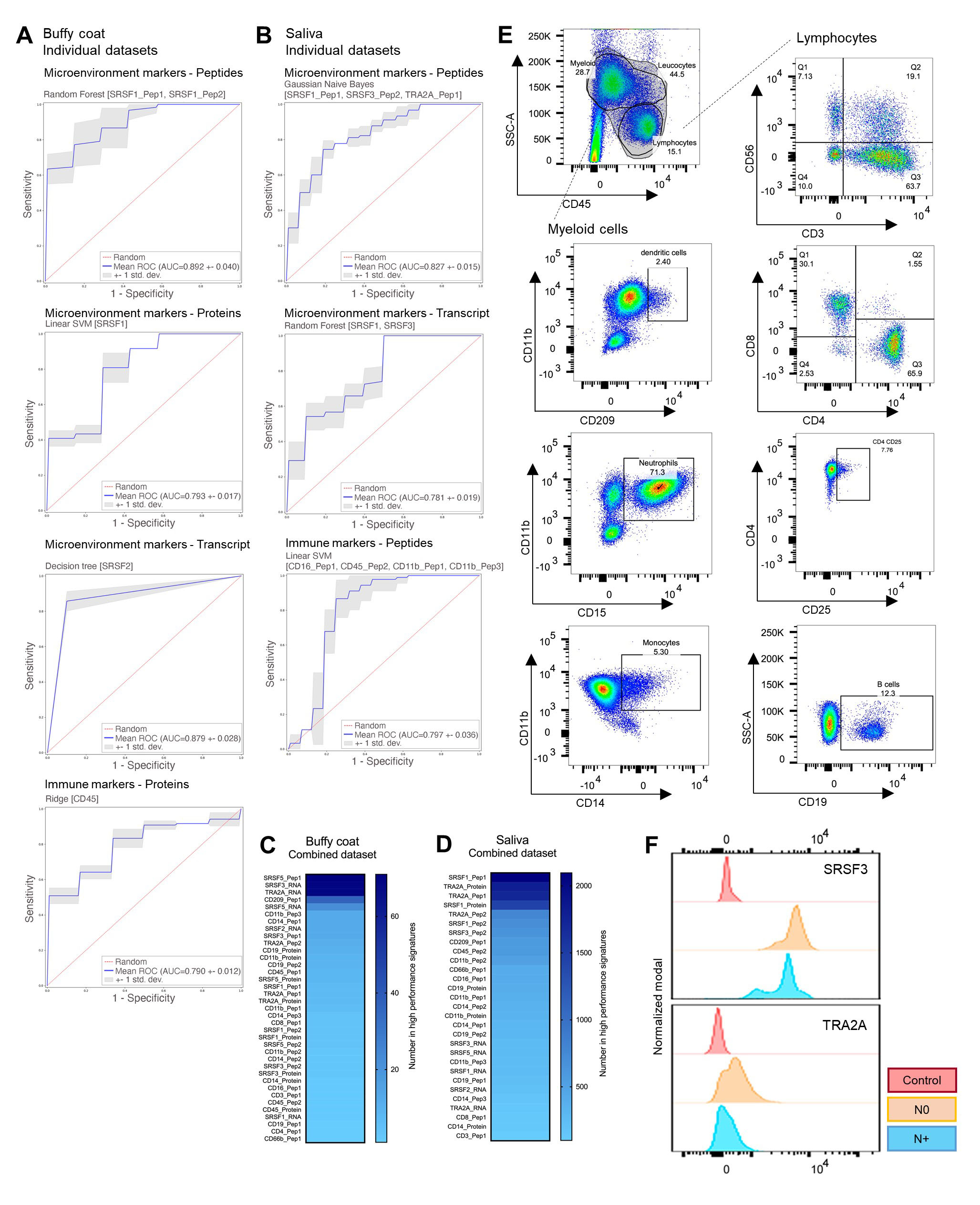
